# ESCRT disruption provides evidence against transsynaptic signaling functions for extracellular vesicles

**DOI:** 10.1101/2023.04.22.537920

**Authors:** Erica C. Dresselhaus, Kathryn P. Harris, Cassandra R. Blanchette, Kate Koles, Steven J. Del Signore, Matthew F. Pescosolido, Biljana Ermanoska, Mark Rozencwaig, Rebecca C. Soslowsky, Michael J. Parisi, Bryan A. Stewart, Timothy J. Mosca, Avital A. Rodal

**Affiliations:** Department of Biology, Brandeis University, Waltham, MA; Office of the Vice-Principal, Research and Innovation, University of Toronto, Mississauga, Mississauga, Canada; Department of Biology, University of Toronto Mississauga, Mississauga, Canada; Department of Cell and Systems Biology University of Toronto, Toronto, Canada; Department of Neuroscience, Vickie and Jack Farber Institute of Neuroscience, Thomas Jefferson University, Bluemle Life Sciences Building, Philadelphia, PA

**Keywords:** ESCRT, Tsg101, Hrs, Shrub, Vps4, Amyloid Precursor Protein, Synaptotagmin-4, Neuroglian, Evi, Wingless, Draper, *Drosophila*, extracellular vesicle, endosome, exosome, synapse

## Abstract

Extracellular vesicles (EVs) are released by many cell types including neurons, carrying cargoes involved in signaling and disease. It is unclear whether EVs promote intercellular signaling or serve primarily to dispose of unwanted materials. We show that loss of multivesicular endosome-generating ESCRT (endosomal sorting complex required for transport) machinery disrupts release of EV cargoes from *Drosophila* motor neurons. Surprisingly, ESCRT depletion does not affect the signaling activities of the EV cargo Synaptotagmin-4 (Syt4) and disrupts only some signaling activities of the EV cargo Evenness Interrupted (Evi). Thus, these cargoes may not require intercellular transfer via EVs, and instead may be conventionally secreted or function cell autonomously in the neuron. We find that EVs are phagocytosed by glia and muscles, and that ESCRT disruption causes compensatory autophagy in presynaptic neurons, suggesting that EVs are one of several redundant mechanisms to remove cargoes from synapses. Our results suggest that synaptic EV release serves primarily as a proteostatic mechanism for certain cargoes.

## Introduction

Neurons release extracellular vesicles (EVs) that can mediate intercellular communication, dispose of unwanted neuronal components, and propagate pathological factors in neurodegenerative disease (Budnik et al., 2016; Holm et al., 2018; Song et al., 2020). Many elegant functional studies of neuronal EVs involve their purification from donor cells and subsequent application to target cells for tests of biological activity e.g. (Gong et al., 2016; Vilcaes et al., 2021). These experiments demonstrate that EVs containing specific cargoes are sufficient to cause functional changes in the recipient cell, but do not rigorously show that traffic into EVs is necessary for the functions of cargoes *in vivo*. In the donor cell, EV cargoes are typically trafficked through the secretory system, plasma membrane, and endosomal network, where they might execute intracellular activities in the donor cell before being released (van Niel et al., 2018). Therefore, to test the physiological functions of EVs *in vivo*, it will be essential to uncouple potential donor cell-autonomous from transcellular functions of these cargoes, using tools that specifically block EV release. Developing such tools will require a deeper understanding of how cargoes are packaged into EVs, and released in a spatially and temporally controlled fashion, especially within the complex morphology of neurons (Blanchette and Rodal, 2020).

Exosomes are a type of EV that arise when multivesicular endosomes (MVEs) fuse with the plasma membrane, releasing their intralumenal vesicles (ILVs) into the extracellular space. Spatial and temporal regulation of the machinery that controls formation of MVEs is therefore likely to be critical for exosome cargo selection, packaging, and release. MVEs can form via multiple nonexclusive mechanisms for budding of vesicles into the endosomal lumen (van Niel et al., 2018). One such pathway relies on Endosomal Sorting Complex Required for Traffic (ESCRT) proteins. In this pathway, ESCRT-0, -I, and -II components cluster cargoes, deform membranes, and then recruit ESCRT-III components, which form a helical polymer that drives fission of the ILV. The VPS4 ATPase remodels and finally disassembles the ESCRT-III filaments (Gruenberg, 2020; Vietri et al., 2020). The ESCRT-I component Tsg101 (Tumor susceptibility gene 101) is incorporated into and serves as a common marker for EVs, highlighting the link between ESCRT and EVs (van Niel et al., 2018). A neutral sphingomyelinase (nSMase)-mediated pathway may operate together with or in parallel to ESCRT to generate EVs by directly modifying lipids and altering their curvature, and indeed EV release of many neuronal cargoes is sensitive to nSMase depletion or inhibition (Asai et al., 2015; Dinkins et al., 2016; Goncalves et al., 2015; Guo et al., 2015; Men et al., 2019; Sackmann et al., 2019). The ESCRT machinery also has functions beyond MVE formation, including autophagosome closure and organelle repair, which are in turn involved in alternative modes of EV biogenesis (Arbo et al., 2020; Lefebvre et al., 2018; Leidal and Debnath, 2021). Further, ESCRT is involved in budding of EVs directly from the plasma membrane (van Niel et al., 2018). However, as there is evidence both for and against a role for ESCRT in EV biogenesis in different neuronal cell types (Cone et al., 2020; Coulter et al., 2018; Gong et al., 2016), it remains unclear whether organism-level physiological defects arising from ESCRT disruption (including ESCRT-linked human neurological disease) could arise from defects in EV traffic (Brugger et al., 2024; Rodger et al., 2020; Sadoul et al., 2018; Ugbode and West, 2021).

At the *Drosophila* larval neuromuscular junction (NMJ), EVs are released from presynaptic motor neurons into extrasynaptic space within the muscle membrane subsynaptic reticulum, and can also be taken up by muscles and glia (Fuentes-Medel et al., 2009; Koles et al., 2012). These EVs are likely to be exosomes, as cargoes are found in presynaptic MVEs, and depend on endosomal sorting machinery for their release and regulation (Blanchette et al., 2022; Koles et al., 2012; Korkut et al., 2009; Lauwers et al., 2018; Walsh et al., 2021).

This system provides the powerful advantage of investigating endogenous or exogenous EV cargoes with known physiological functions in their normal tissue and developmental context. EV cargoes characterized to date at the *Drosophila* NMJ include Synaptotagmin-4 (Syt4, which mediates functional and structural plasticity), Amyloid Precursor Protein (APP, a signaling protein involved in Alzheimer’s Disease), Evenness Interrupted/Wntless/Sprinter (Evi, which carries Wnt/Wingless (Wg) to regulate synaptic development and plasticity), and Neuroglian (Nrg, a cell adhesion molecule) (Koles et al., 2012; Korkut et al., 2013; Walsh et al., 2021). Mutants in membrane trafficking machinery (e.g. *evi*, *rab11 (*a recycling endosome GTPase)), and *nwk* (a component of the endocytic machinery) cause reduced levels of cargo in EVs and show defects in the physiological activities of EV cargoes (Blanchette et al., 2022; Korkut et al., 2009; Korkut et al., 2013), leading to the hypothesis that trans-synaptic transfer of these cargoes into the postsynaptic muscle is required for their signaling functions (Budnik et al., 2016). However, we and others have shown that these mutants also have a dramatic local presynaptic decrease in cargo levels, making it difficult to rule the donor neuron out as their site of action (Ashley et al., 2018; Blanchette et al., 2022; Koles et al., 2012; Korkut et al., 2009; Korkut et al., 2013; Walsh et al., 2021). Here we show that disruption of the ESCRT machinery can cause a specific loss of EV release without strongly depleting presynaptic cargo levels, and use this system to test whether EV release is required for cargo signaling functions.

## Results

### ESCRT machinery promotes EV release from synapses

To determine if the ESCRT pathway is involved in EV release at the *Drosophila* NMJ synapse, we first used GAL4/UAS-driven RNAi to knock down the ESCRT-I component Tsg101 (Tumor susceptibility gene 101) specifically in neurons (Tsg101^KD^). We then used our previously established methods to measure the levels of the endogenously tagged EV cargo Syt4-GFP, both in the donor presynaptic compartment and in neuron-derived EVs in the adjacent postsynaptic cleft and muscle (Walsh et al., 2021). Neuronal knockdown of Tsg101 (Tsg101^KD^) led to accumulation of Syt4-GFP in the presynaptic compartment, together with a striking loss of postsynaptic Syt4-GFP EVs (**Fig 1A, E**). We next tested the effects of Tsg101^KD^ on three other known EV cargoes: neuronal UAS-driven Evi-GFP **(1B, F)** or human APP-GFP **(1C, G)**, and endogenous Nrg **(1D, H)** (Korkut et al., 2009; Korkut et al., 2013; Walsh et al., 2021). For all three cargoes, we observed a similar phenotype to Syt4-GFP: presynaptic redistribution in large structures (accumulating to particularly high levels for Syt4-GFP and Evi-GFP), together with loss of postsynaptic EV signal. Similarly, neuronal UAS-driven Tsp42Ej/Sunglasses (a *Drosophila* tetraspanin EV marker (Walsh et al., 2021)) was strongly depleted from the postsynaptic region upon Tsg101^KD^ (**Fig. S1A**).

**Fig 1.**
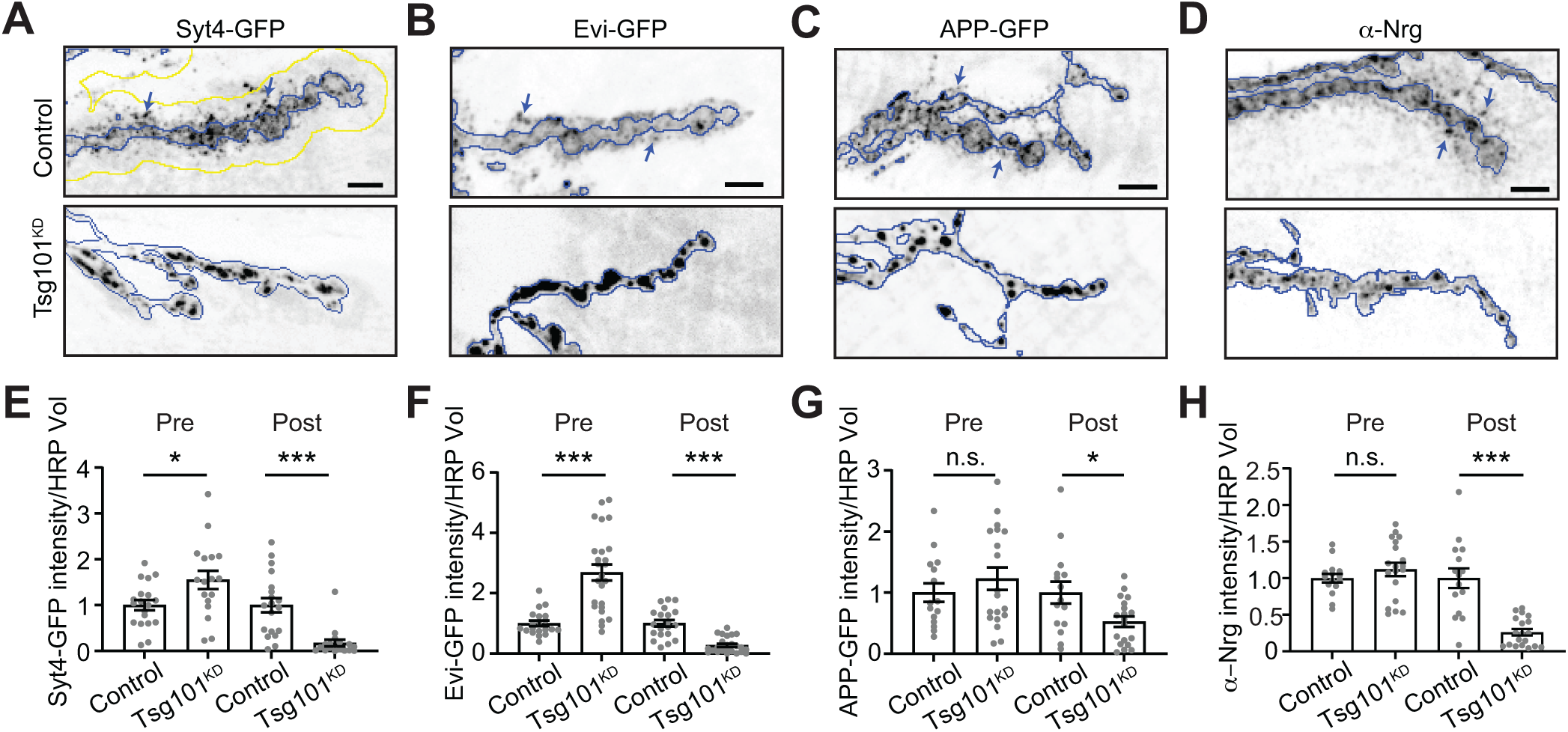
Tsg101 is required for release of EV cargoes from presynaptic terminals. **(A-D)**Representative confocal images from larvae expressing UAS-Tsg101-RNAi (Tsg101^KD^) or a control RNAi either pan-neuronally (C380-GAL4) or in motor neurons (Vglut-GAL4) together with the following EV cargoes: **(A)** Syt4-GFP expressed from its endogenous locus, **(B)** UAS-driven Evi-GFP, **(C)** UAS-driven APP-GFP, **(D)** endogenous Neuroglian (Nrg, neuronal isoform Nrg180) detected by antibody. **(E-H)** Quantification of EV cargo puncta intensity. All images show MaxIPs of muscle 6/7 segments A2 or A3. Scale bars are 5 µm. **(A-D)** Blue outlines represent the neuronal membrane as marked from an HRP mask; yellow line in **(A)** shows a 3.3 µm dilation of the HRP mask, representing the postsynaptic region. Arrows show examples of postsynaptic EVs. Data are represented as mean +/-s.e.m.; n represents NMJs. All intensity measurements are normalized to their respective controls. *p<0.05, ***p<0.001. **See Tables S1 and S3 for detailed genotypes and statistical analyses.**

We then used super-resolution STED microscopy to measure the size and number of presynaptically-derived EV cargoes in the 3 µm region surrounding the presynaptic terminal. The density of postsynaptic puncta was strongly reduced at Tsg101^KD^ synapses (**Fig. S1B-E**), and was not significantly different from background signal. Further, the diameters of Nrg and APP-GFP extraneuronal puncta were ∼125 nm, consistent with the size of MVE-derived exosomes (Welsh et al., 2024)). Given their size, the observation that cargoes accumulate in internal structures upon Tsg101^KD^, and previous observations of MVE fusion with the plasma membrane at this synapse (Koles et al., 2012; Korkut et al., 2009; Lauwers et al., 2018), it is most likely that NMJ EVs are exosomes derived from intracellular MVEs, rather than budding directly from the plasma membrane. Thus, multiple EV cargoes, either endogenously or exogenously expressed, require the ESCRT-I component Tsg101 for release in neuronally-derived EVs.

In addition to its functions in MVE biogenesis, Tsg101 also plays roles in numerous cellular processes including membrane repair, lipid transfer, neurite pruning, and autophagy, each depending on a specific subset of other ESCRT machinery (Vietri et al., 2020). We therefore tested if EV release depends on other canonical ESCRT components. Hrs (Hepatocyte growth factor receptor substrate) is a component of the ESCRT-0 complex and is required to cluster EV cargo on the delimiting membrane of the endosome (Vietri et al., 2020). Similar to Tsg101^KD^, *Hrs* loss-of-function mutants caused a strong decrease in postsynaptic Syt4-GFP, Evi-GFP, and Nrg, though interestingly their presynaptic levels were also partially depleted, unlike the Tsg101^KD^ condition (**Fig 2A-F**). Notably, direct comparison showed that *Hrs* mutants exhibited a postsynaptic decrease in Syt4-GFP nearly as severe as *nwk* mutants, but with a comparatively modest decrease in presynaptic Syt4-GFP (**Fig 2A,D**).

**Fig 2.**
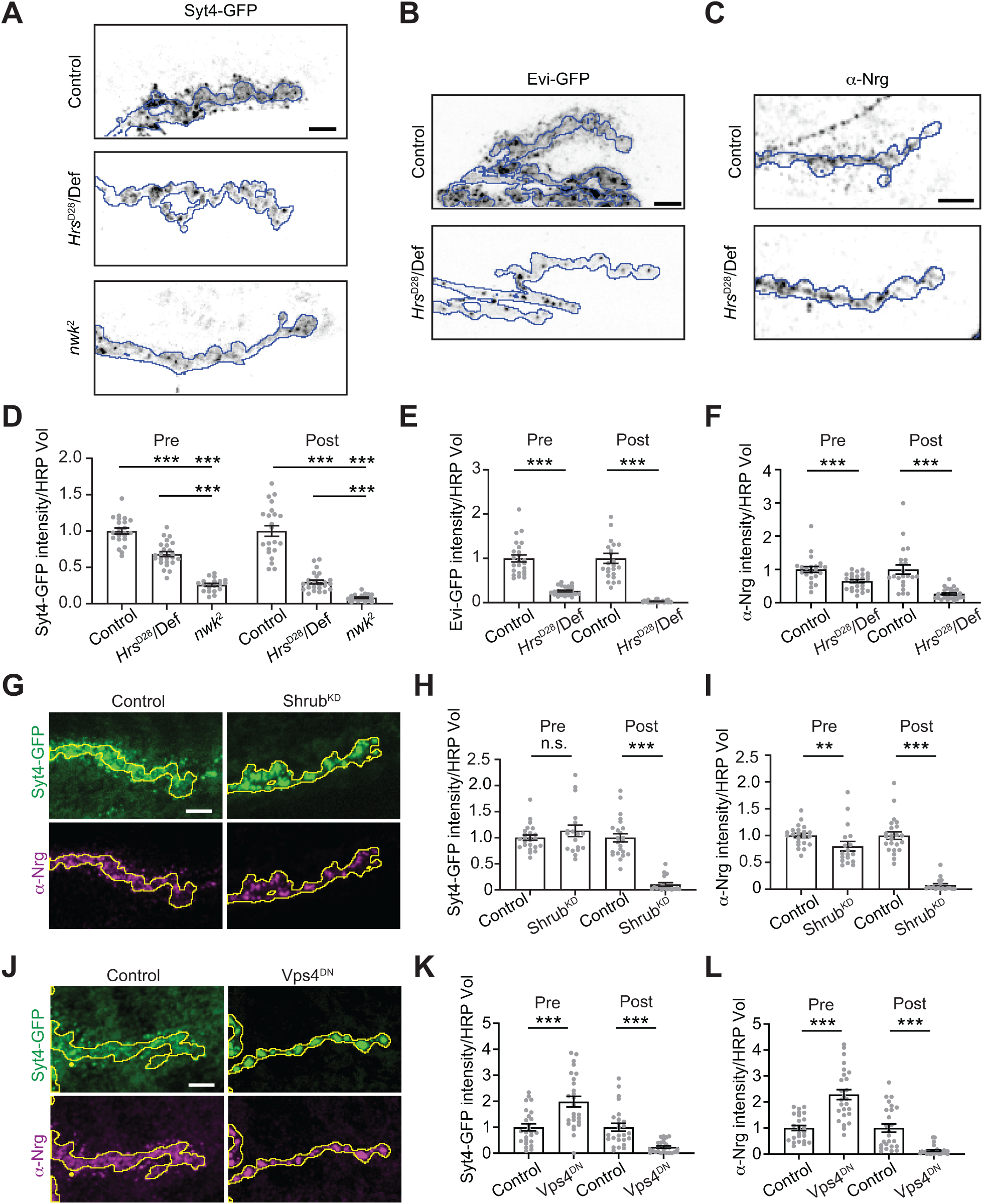
Multiple ESCRT components are required for release of EV cargoes from presynaptic terminals. **(A)** Representative confocal images of control, *Hrs* and *nwk* mutant larvae expressing Syt4-GFP from its endogenous locus. **(B)** Representative confocal images of control and *Hrs* mutant larvae expressing motor neuron (Vglut-GAL4)-driven UAS-Evi-GFP. **(C)** Representative confocal images of control and *Hrs* mutant larvae labeled with antibodies against endogenous Nrg. **(D-F)** Quantification of EV cargo puncta intensity. **(G)** Representative confocal images of larvae pan-neuronally expressing UAS-Shrub-RNAi (Shrub^KD^) or a control RNAi. **(H-I)** Quantification of Syt4-GFP and Nrg puncta intensity. **(J)** Representative confocal images of larvae pan-neuronally expressing UAS-Vps4^DN^. **(K-L)** Quantification of Syt4-GFP and Nrg puncta intensity. All images show MaxIPs of muscle 6/7 segments A2 or A3. Scale bars are 5 µm. Outline represents the neuronal membrane as marked from an HRP mask. Data are represented as mean +/- s.e.m.; n represents NMJs. All fluorescence intensity values are normalized to their respective controls. *p<0.05, **p<0.01, ***p<0.001. **See Tables S1 and S3 for detailed genotypes and statistical analyses.**

Next, we tested ESCRT-III, which forms the polymer involved in constriction and scission of the ILV neck. The *Drosophila* genome encodes several ESCRT-III proteins, of which *shrub* is homologous to mammalian CHMP4B. Shrub is likely to play an important role at synapses, since its loss leads to defects in NMJ morphogenesis and ILV formation (Sweeney et al., 2006). Pan-neuronal RNAi of *shrub* (Shrub^KD^) caused a dramatic loss of postsynaptic Syt4-GFP and Nrg signals (**Fig. 2G-I**). Finally, we examined the role of Vps4, which catalyzes remodeling and disassembly of the ESCRT-III polymer, finalizing the formation of the ILV. Pan-neuronal expression of a dominant negative Vps4 fragment (Vps4^DN^, (Rodahl et al., 2009)) strongly reduced postsynaptic levels of both Syt4-GFP and Nrg, and increased their presynaptic levels (**Fig. 2J-L**). Together, these results demonstrate that multiple components of the ESCRT pathway are required for release of EV cargoes at neuronal synapses, with variable effects on presynaptic accumulation of these cargoes.

### Loss of Tsg101 or Hrs uncouples autophagic and EV functions of ESCRT machinery

To explore the nature of the presynaptic accumulations of EV cargoes at Tsg101^KD^ NMJs, we examined their co-localization with early (Rab5) and recycling (Rab11) endosomes, which drive an endosome-to-plasma membrane recycling flux that supplies the EV biogenesis pathway at this synapse (Korkut et al., 2009; Walsh et al., 2021). We also examined cargo co-localization with late endosomes (Rab7), which play less important roles in NMJ EV traffic (Walsh et al., 2021). We found that EV cargoes exhibited increased co-localization with all these endosomal markers at Tsg101^KD^ synapses, in what appeared to be multi-endosome clusters (**Fig 3A, S2A-C**). These results argue against formation of a single type of arrested MVE such as the canonical Class E compartment in ESCRT-deficient yeast and mammalian cells (Doyotte et al., 2005; Raymond et al., 1992) and instead suggest a more global defect in endosome maturation or turnover. To test this hypothesis, we measured the overall mean intensity as well as puncta number and size for Rab5, Rab7, and Rab11 upon Tsg101 knockdown. We saw no changes in the total intensity or puncta parameters for Rab7 (**Fig S2D-F**). However, for both Rab5 and Rab11, we observed a significant increase in Rab puncta intensity, mean intensity over the whole NMJ, and an increase in puncta size, with no change (Rab11) or a slight decrease (Rab5) in the number of puncta (**Fig S2D-F)**. These results suggest defects in early and recycling endosome maturation and/or turnover upon loss of ESCRT function.

**Figure 3:**
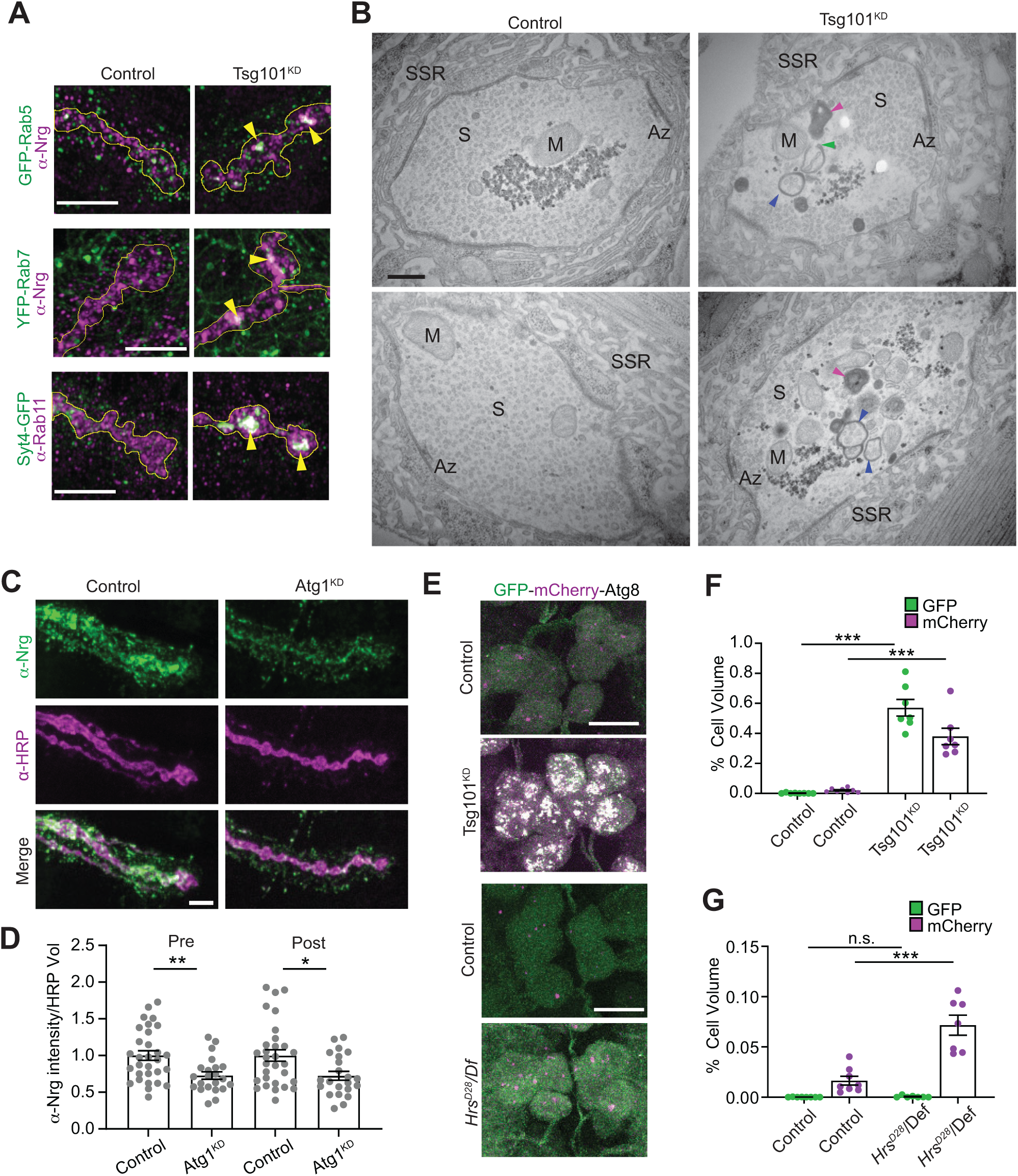
Loss of ESCRT causes compensatory autophagy of presynaptic cargoes. **(A)** Representative Airyscan images showing co-localization of EV cargoes Syt4-GFP or α-Nrg with endosomal markers α-Rab11, GFP-Rab5 (endogenous tag), or YFP-Rab7 (endogenous tag). Scale bars are 5 μm and outline represents the neuronal membrane as marked from an HRP mask. **(B)** Representative TEM images of boutons from muscle 6/7 from wild-type and neuronal Tsg101^KD^ larvae. Examples of autophagic vacuoles are marked with arrowheads, blue = autophagosome, magenta = autolysosome, and green = unclosed phagophore. Other notable features include Az = active zone, S = synaptic vesicles, M = mitochondria, SSR = subsynaptic reticulum. Scale bar is 400 nm. **(C)** Representative images of the EV cargo Nrg following motor neuron knockdown of *Atg1*. Scale bar is 5 μm. **(D)** Quantification of Nrg intensity from **(C**), normalized to control. **(E)** Representative images from neuronal cell bodies in the ventral ganglion expressing motor neuron-driven GFP-mCherry-Atg8. Scale bar is 10 μm. Brightness/contrast are matched for each mutant with its paired control (**see Table S3**). **(F-G)** Quantification of GFP-mCherry-Atg8 levels in **(F)** Tsg101^KD^ and **(G)** *Hrs^D28^* mutant larvae. Data are represented as mean +/- s.e.m.; n represents NMJs in **(C)** and animals in **(F-G**). *p<0.05, **p<0.01, ***p<0.001. **See Tables S1 and S3 for detailed genotypes and statistical analyses.**

We next examined Tsg101^KD^ NMJs using transmission electron microscopy (TEM). Tsg101^KD^ boutons retained typical mitochondria, synaptic vesicles and active zone “T-bars”, and were surrounded by an apparently normal subsynaptic reticulum, representing the infolded postsynaptic muscle membrane. However, within Tsg101^KD^ boutons we observed striking clusters of double membrane-surrounded or electron-dense structures, typical of autophagic vacuoles at various stages of maturation (Klionsky et al., 2021), including those with unclosed phagophores (**Fig. 3B**; three or more autophagic vacuoles were observed in 58.9% of mutant boutons (n=56) compared to 2.5% of control boutons (n=40), p<0.001). Given that secretion of autophagosomal contents has been described as an EV-generating mechanism (Buratta et al., 2020), we next tested whether core autophagy machinery might play a role in EV release at the *Drosophila* NMJ, and could therefore be linked to the Tsg101^KD^ EV phenotype. Atg1 is a kinase that is required for the initiation of phagophore assembly, and acts as a scaffold for recruitment of subsequent proteins, while Atg2 is required for phospholipid transfer to the phagophore (Nakatogawa, 2020). We observed a modest but significant decrease in both pre- and post-synaptic levels of the EV marker Nrg upon disruption of autophagy by RNAi-mediated knockdown of Atg1 (knockdown validated in **Fig. S2G**, Nrg results in **Fig. 3C-D**), as well as by loss-of-function *Atg2* mutations (**Fig. S2H-I**). Notably, these mutants did not recapitulate the ESCRT mutant phenotype of dramatic depletion of postsynaptic EVs and presynaptically trapped cargoes. These results indicate that autophagic machinery does not play a major role in EV biogenesis or release at the NMJ, and that the autophagic defects at Tsg101^KD^ NMJs are likely separable from its roles in EV release.

To further explore these autophagic defects, we next used the reporter GFP-mCherry-Atg8/LC3 to assess autophagic flux in Tsg101^KD^ neurons. Under normal circumstances, the GFP moiety in this reporter is quenched when autophagosomes fuse with acidic endosomes and lysosomes, while mCherry retains its fluorescence. By contrast, defects in autophagic flux lead to accumulation of structures with both GFP and mCherry fluorescence (Klionsky et al., 2021). We examined this flux in motor neuron cell bodies, where mature autolysosomes are predicted to accumulate (Sidibe et al., 2022). In wild-type animals we observed mCherry-positive/GFP-negative puncta reflecting mature autolysosomes. By contrast we observed an increased volume of intense puncta in the cell bodies of Tsg101^KD^ motor neurons, most of which were labeled by both mCherry and GFP (**Fig. 3E-F, S2J**). These data suggest that Tsg101^KD^ causes a defect in autophagic flux.

Since autophagy is normally rare at wild-type *Drosophila* NMJ synapses (Soukup et al., 2016), we hypothesized that ESCRT mutants might activate a compensatory “endosomophagy” or “simaphagy” pathway to degrade ESCRT-deficient endosomes (Migliano et al., 2023; Millarte et al., 2022). However, since Tsg101 is also required for phagophore closure (Takahashi et al., 2018), this process is likely unable to dispose of defective endosomes upon Tsg101 knockdown. By contrast, ESCRT-0/Hrs is not required for autophagy in some cell types, such as *Drosophila* fat body (Rusten et al., 2007). To test if this also applies to motor neurons, we examined GFP-mCherry-Atg8 in *Hrs* mutant motor neuron cell bodies, and found mCherry-positive, GFP-negative structures, similar to controls (**Fig. 3E, G, S3J**), suggesting that Hrs is not required for autophagic flux in motor neurons. Interestingly, we did observe an increase in the area covered by puncta, indicating that autophagy is induced in *Hrs* mutants (**Fig. 3G**). Overall, *Tsg101* and *Hrs* have different autophagy phenotypes but similar EV release defects, and canonical autophagy mutants do not phenocopy ESCRT mutants in trapping EV cargoes presynaptically. Together, these results suggest that autophagy and EV traffic are separable functions of ESCRT at the synapse, and that a compensatory (and Tsg101-dependent) autophagy mechanism might be activated to remove defective endosomes in *Hrs* mutants.

Finally, we further explored whether accumulation of EV cargoes in arrested structures was local to the synapse or occurring throughout the neuron. First, we examined Syt4-GFP levels in motor neuron cell bodies and axons after Tsg101 knockdown, and found that Syt4-GFP accumulated significantly at both locations (**Fig. 4A-B**). To ask whether the presynaptic accumulations could be due to faster anterograde or slowed retrograde transport of EV cargo-containing compartments, we next conducted live imaging and kymograph analysis of motor neuron-driven APP-GFP, as well as a mitochondrial marker. We found that Tsg101 knockdown led to a large increase in the number of stationary APP-GFP puncta in axons without affecting the number of compartments undergoing retrograde or anterograde transport (**Fig 4C-D**), though we observed a small decrease in the retrograde transport rate (**Fig 4E**). By contrast, we did not observe an increase in the steady state intensity of the mitochondrial marker or see any changes in its transport behavior (**Fig S3A-D**), suggesting that axonal accumulations are specific to EV cargo. Thus, loss of *tsg101* leads to accumulation of stationary EV cargo-containing compartments throughout the neuron, without affecting the transport rates of moving cargoes, suggesting that altered axonal transport kinetics do not underlie synaptic accumulation.

**Figure 4.**
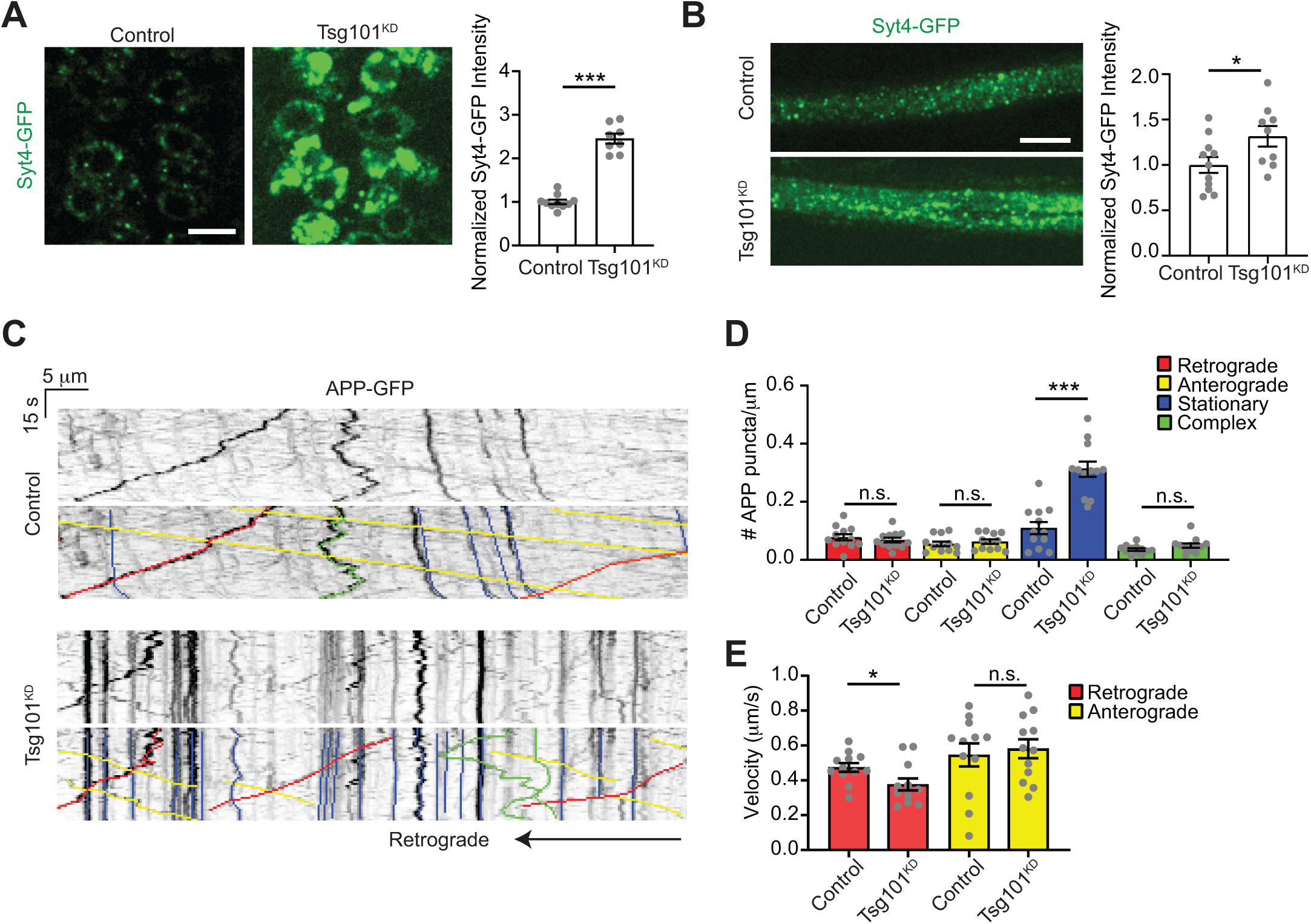
Tsg101^KD^ causes neuronal accumulation of EV cargoes. **(A) (Left)** Representative confocal images of Syt4-GFP in a single slice through motor neuron cell bodies of the ventral ganglion. Scale bar is 10 μm. **(Right)** Quantification of total Syt4-GFP intensity in the brain. **(B) (Left)** Maximum intensity projection of axon segment proximal to the ventral ganglion. Scale bar is 10 μm. **(Right)** Quantification of total Syt4-GFP intensity in the axon. **(C)** Representative kymographs showing tracks of APP-GFP in the axon proximal to the ventral ganglion. Bottom panels show color coded traces. **(D)** Quantification of directionality of APP-GFP tracks. **(E)** Quantification of the velocity of retrograde and anterograde APP-GFP tracks upon neuronal Tsg101^KD^. Data are represented as mean +/- s.e.m.; n represents animals. Intensity measurements (**A, B**) are normalized to their respective controls. *p<0.05, ***p<0.001. **See Tables S1 and S3 for detailed genotypes and statistical analyses.**

### *evi* and *wg* signaling are not correlated with EV release

Specific depletion of cargo in postsynaptic EVs (but not the donor presynaptic terminal) upon ESCRT disruption provided us with a valuable tool to determine if these cargoes require trans-synaptic transfer for their known synaptic functions. Neuron-derived Wg provides anterograde (to the muscle) and autocrine (to the neuron) signals, promoting NMJ growth, active zone development, and assembly of the postsynaptic apparatus (Miech et al., 2008; Packard et al., 2002). Evi is a multipass transmembrane protein that serves as a carrier for Wg through the secretory system, ultimately leading to Wg release from the cell, either by conventional exocytosis or via EVs (Das et al., 2012). At the NMJ, Evi co-transports with Wg into EVs, and *evi* mutants phenocopy *wg* signaling defects, providing support for the hypothesis that Evi/Wg EVs are required for Wg signaling (Koles et al., 2012; Korkut et al., 2009). However, since Evi is broadly required for many steps of Wg traffic, *evi* mutants trap Wg in the somatodendritic compartment and prevent its transport into presynaptic terminals (Korkut et al., 2009). Therefore, Wg signaling defects in *evi* mutants may be due to generalized loss of Wg secretion rather than specific loss of its trans-synaptic transfer. *Wg* or *evi* mutants exhibit dramatic reductions in bouton number, together with the appearance of immature boutons with abnormal or missing active zones, fewer mitochondria, aberrant swellings or pockets in the postsynaptic region opposing active zones, and missing areas of PSD95/Discs-Large (DLG)-positive postsynaptic subsynaptic reticulum (Korkut et al., 2009; Packard et al., 2002). We found that in *evi* mutants, the number of synaptic boutons and the number of active zones (marked by ELKS/CAST/ Bruchpilot (BRP)) were both significantly reduced compared to controls, and the postsynaptic scaffolding molecule DLG frequently exhibited a “feathery” appearance, suggesting defects in postsynaptic assembly (**Fig. 5A-E**). By contrast, we found that bouton and active zone numbers at Tsg101^KD^ NMJs (which have presynaptic Evi but strongly diminished Evi EVs (**Fig. 1B**)) were not significantly different from controls. Further, active zones in Tsg101^KD^ appeared morphologically normal by TEM (**Fig. 3B**). We did not observe a significant frequency of “feathery” DLG distribution in control or Tsg101^KD^ larvae (2.3% of control NMJs (n=87) and 5.5% of Tsg101^KD^ NMJs (n=87), compared to 51.7% of *evi*^2^ mutant NMJs (n=91), p=0.27 for control versus Tsg101^KD^). We also did not observe significant differences between control and Tsg101^KD^ NMJs in the appearance of subsynaptic reticulum by EM (**Fig. 3B**) These results indicate that some neuronal Evi and Wg functions are unexpectedly maintained despite loss of detectable postsynaptic Evi EVs upon Tsg101^KD^.

**Figure 5.**
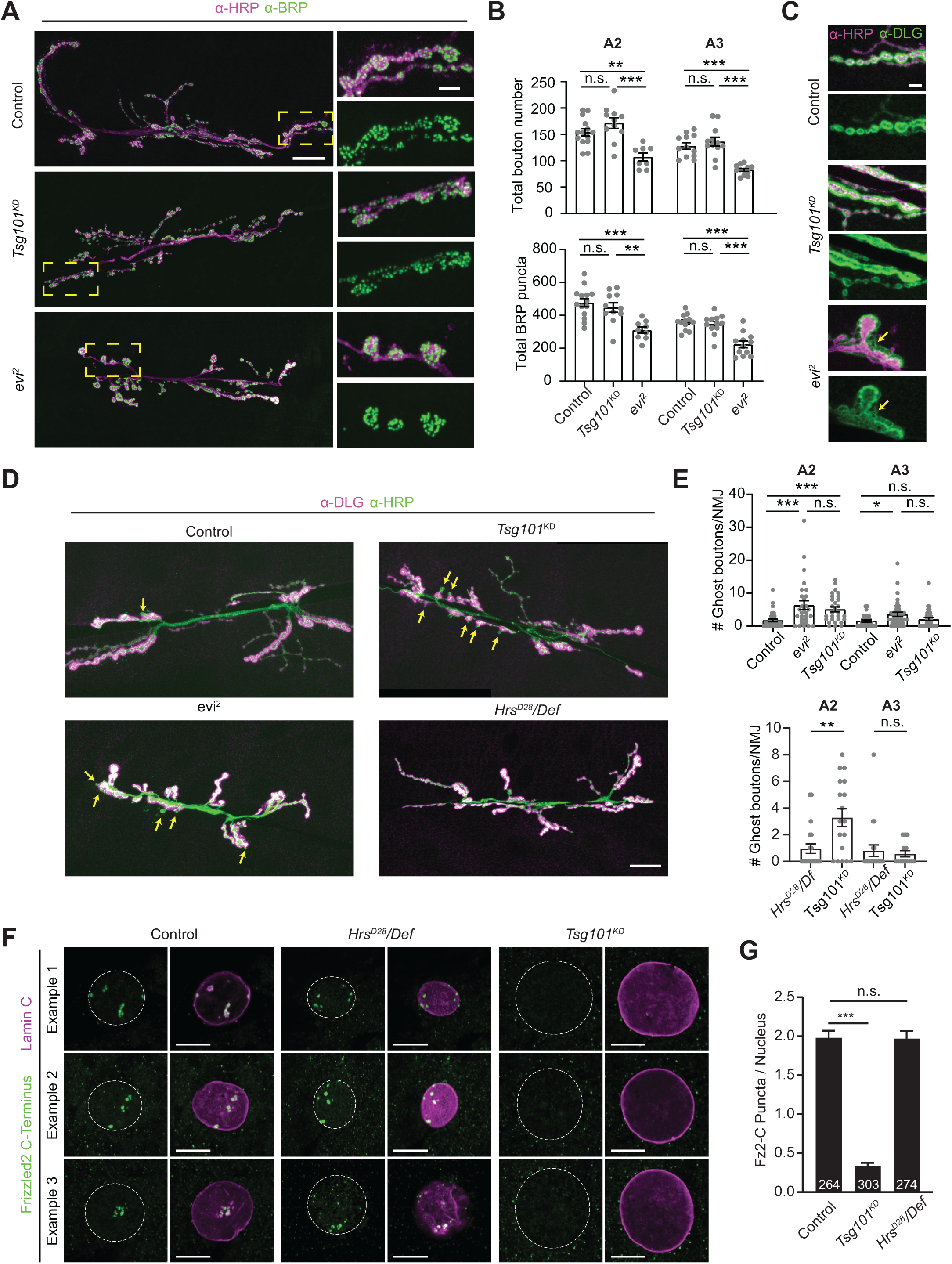
Tsg101^KD^ phenocopies a subset of *evi* and *wg* synaptic morphology and signaling defects, while loss of Hrs has no effect. **(A)** Representative confocal images of muscle 6/7 NMJs labeled with α-HRP and α-BRP antibodies (left). Magnification of the yellow boxed area (right). HRP brightness was adjusted independently. Large image scale bar is 20 µm, small image scale bar is 5 µm. **(B)** Quantification of total bouton number (top) and active zone number (bottom) on muscle 6/7. **(C)** Representative confocal images of muscle 6/7 NMJ highlighting α-DLG pattern. Arrows indicate location of “feathery” DLG. Scale bar is 5 µm. **(D)** Representative confocal images of muscle 6/7 NMJ (abdominal segment A2) labeled with α-HRP and α-DLG antibodies. α-DLG and α-HRP signals were acquired in the linear range but adjusted independently and displayed near saturation to highlight DLG-negative ghost boutons, which are indicated with yellow arrows. **(E)** Quantification of baseline (i.e. unstimulated) ghost boutons. Top and bottom graphs represent independent experiments. **(F)** Single slices of muscle 6/7 nuclei labeled with α-LamC (nuclear periphery) and α-Fz2-C antibodies. Dotted line represents LamC-defined nuclear boundary. Scale bars are 10µm. **(G)** Quantification of Fz2-C puncta per nucleus. Number of nuclei quantified are indicated in the bar graph. A2 and A3 indicate the larval abdominal segment. Data are represented as mean +/- s.e.m.; n represents nuclei in **(G)** and NMJs in (**B-E**). *p<0.05, **p<0.01, ***p<0.001. **See Tables S1 and S3 for detailed genotypes and statistical analyses.**

In addition to an overall decrease in the number of synaptic boutons, both *Wg* and *evi^2^* mutants show increased numbers of developmentally arrested or “ghost” boutons that feature presynaptic markers such as α-HRP antigens, but lack a postsynaptic apparatus defined by DLG (Korkut et al., 2009). We found that in *evi*^2^ mutants, these ghost boutons are more prevalent in anterior segments of the larvae, where overall synaptic growth is more exuberant. Similarly, Tsg101^KD^ animals exhibited a significant increase in ghost boutons in abdominal segment A2 (but not in A3), partially phenocopying the *evi*^2^ mutant (**Fig, 5 D,E**). In sharp contrast, *Hrs* mutants did not exhibit a significant change in ghost bouton number despite having a similar decrease in postsynaptic Evi-GFP to Tsg101^KD^ (**Fig, 5 D,E**). These results suggest that Evi release in EVs and *wg*-related phenotypes can be uncoupled.

To further explore Wg signaling in ESCRT mutants, we directly measured the output of this pathway. In *Drosophila* muscles, Wg does not signal via the conventional β-catenin pathway. Instead, neuronally-derived Wg activates cleavage of its receptor Fz2, resulting in translocation of a Fz2 C-terminal fragment into muscle nuclei (Mathew et al., 2005; Mosca and Schwarz, 2010). Using an antibody specific to the Fz2 C-terminus, we measured the number of nuclear Fz2-C puncta (**Fig. 5F-G**). *Hrs* mutants showed a similar number of puncta compared to controls. By contrast neuronal knockdown of Tsg101 caused a dramatic loss of Fz2-C puncta, consistent with our findings for ghost boutons. Given the similarly strong loss of EVs in *Hrs* mutants and Tsg101^KD^, these results indicate that EVs are not required for transynaptic signaling by Wg, and suggest that a separable membrane trafficking pathway for Wg secretion is defective only in the Tsg101^KD^ condition.

### ESCRT loss does not recapitulate *syt4* phenotypes in activity-dependent structural or functional plasticity

We next explored the functions of the EV cargo Syt4, which is required for activity-dependent structural and functional plasticity at the NMJ (Barber et al., 2009; Korkut et al., 2013; Piccioli and Littleton, 2014; Yoshihara et al., 2005). Endogenous Syt4 is thought to be generated only by the presynaptic motor neuron, based on the absence of Syt4 transcript in muscle preparations, and the finding that presynaptic RNAi of Syt4 depletes both presynaptic and postsynaptic signals (Korkut et al., 2013). We independently verified that all the Syt4 at the NMJ was derived from the neuron, using a strain in which the endogenous Syt4 locus is tagged at its 3’ end with a switchable TagRFP-T tag, which could be converted to GFP in the genome via tissue-specific GAL4/UAS expression of the Rippase recombinase (Koles et al., 2015; Walsh et al., 2021) (**Fig. S4A**). Conversion of the tag only in neurons resulted in a bright Syt4-GFP signal both presynaptically and postsynaptically, together with disappearance of the TagRFP-T signal. However, conversion of the tag in muscles did not result in any GFP signal, and the TagRFP-T signal remained intact (**Fig. S4B**). These results indicate that Syt4 is only expressed in neurons, and support the previous conclusion that the postsynaptic signal is derived from a presynaptically-expressed product (Korkut et al., 2013).

Membrane trafficking mutants such as *rab11* and *nwk* deplete Syt4 from presynaptic terminals, secondarily reducing its traffic into EVs, and phenocopy *syt4* null mutant plasticity phenotypes (Blanchette et al., 2022; Korkut et al., 2013; Walsh et al., 2021). This does not provide conclusive evidence that signaling by Syt4 explicitly requires its transfer via EVs, since Syt4 is missing from both the donor and recipient compartment and could therefore be signaling in the presynaptic cell. Given that loss of *Hrs* and *tsg101* leads to a similar postsynaptic decrease in Syt4 to *nwk* mutants but without a strong presynaptic decrease, these mutants present an opportunity to challenge the hypothesis that Syt4 must transfer via EVs to exert its functions. We first tested the effect of Tsg101^KD^, which depletes the majority of EVs without diminishing presynaptic Syt4 (**Fig. 1A, E**), on Syt4-dependent structural plasticity. In this paradigm, spaced high potassium stimulation promotes acute formation of nascent ghost boutons (Ataman et al., 2006; Korkut et al., 2013; Piccioli and Littleton, 2014). These are likely transient structures, and thus are not directly comparable to developmentally arrested ghost boutons such as those found in *evi* mutants (Fernandes et al., 2023). However, to avoid the confounding presence of these immature boutons, we explored the activity-dependent synaptic growth paradigm on muscle 4, where the Tsg101^KD^ animals do not have significantly more ghost boutons than controls under baseline conditions. Unexpectedly, Tsg101^KD^ animals behaved similarly to controls, and exhibited a significant increase in ghost boutons following high K+ spaced stimulation compared to mock stimulation (**Fig. 6A-B**), suggesting that Syt4 function is preserved in these synapses despite depletion of EV Syt4. We were surprised by these results and contacted another laboratory (KPH, BAS) to replicate this experiment independently at muscle 6/7 in segments A3 and A4, and again readily observed activity-dependent ghost bouton formation (**Fig. S4C-D)**. KPH next tested the effect of Tsg101^KD^ on Syt4-dependent functional plasticity. In this paradigm, stimulation with 4×100 Hz pulses causes a Syt4-dependent increase in the frequency of miniature excitatory junction potentials (mEJPs), in a phenomenon termed High Frequency-Induced Miniature Release (HFMR) (Korkut et al., 2013; Yoshihara et al., 2005).

**Figure 6.**
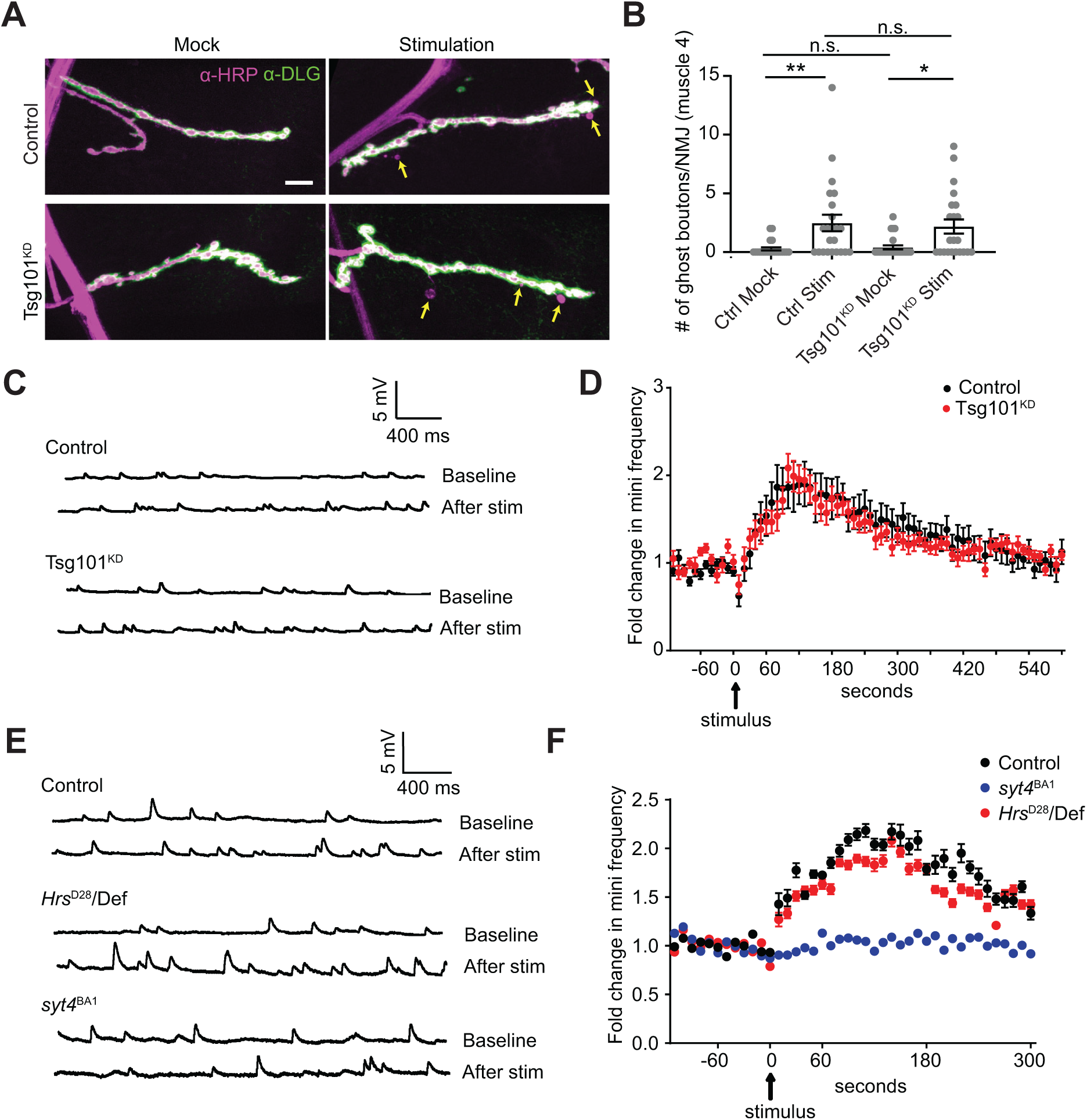
Loss of ESCRT does not phenocopy *syt4* functional defects,. **A)** Representative confocal images from muscle 4 in mock and spaced K^+^-stimulated larvae. Arrows indicate examples of activity-dependent ghost boutons. Scale bar = 10 µm. **(B)** Quantification of ghost bouton numbers per NMJ. **(C)** Representative traces of mEJPs before (top trace) and after (bottom trace) high frequency stimulation (4 × 100 Hz) from control and Tsg101^KD^. **(D)** Timecourse of mEJP frequency after stimulation. **(E)** Representative traces of mEJPs before (top trace) and after (bottom trace) high frequency stimulation (4 × 100 Hz) from control, *Hrs*^D28^, and *syt4*^BA1^. **(F)** Timecourse of mEJP frequency after stimulation. Data are represented as mean +/- s.e.m.; n represents NMJs. *p<0.05, **p<0.01. **See Tables S1 and S3 for detailed genotypes and statistical analyses.**

Tsg101^KD^ animals exhibited similar HFMR to wild type controls, indicating that Syt4 function was not disrupted (**Fig. 6C-D**). *Hrs* mutants, despite being very sickly, also exhibited similar HFMR to wild type controls, in sharp contrast to *syt4* null animals which did not exhibit any HFMR (**Fig. 6E-F**). Taken together, our results show that Syt4-dependent structural and functional plasticity at the larval NMJ can occur despite dramatic depletion of EVs containing Syt4.

### Syt4 is not detectable in the muscle cytoplasm and is taken up by phagocytosis

If trans-synaptic transfer of Syt4 in EVs serves a calcium-responsive signaling function in the muscle, one would expect to find neuronally-derived Syt4 in the muscle cytoplasm.

Therefore, we tested whether neuronally-derived Syt4-GFP (for which the GFP moiety is topologically maintained on the cytoplasmic side of membranes in both donor and recipient cells) could be found in the muscle cytoplasm. Using the GAL4/UAS system, we expressed a proteasome-targeted anti-GFP nanobody (deGradFP (Caussinus et al., 2011), **Fig. 7A**) only in neurons or only in muscles. We observed strong depletion of Syt4-GFP fluorescence upon presynaptic deGradFP expression, including a reduction in Syt4 postsynaptic puncta intensity and number, consistent with the presynaptic source of EV Syt4-GFP protein(**Fig. 7B, D, F**). This result also demonstrates the effectiveness of deGradFP in depleting Syt4-GFP. However, we did not observe any difference in either presynaptic or postsynaptic Syt4-GFP levels or puncta number upon deGradFP expression in the muscle (**Fig. 7C, E, F**), though deGradFP could efficiently deplete DLG as a control postsynaptic protein (**Fig. S5A**). These results suggest that the majority of postsynaptic Syt4 is not exposed to the muscle cytoplasm.

**Figure 7.**
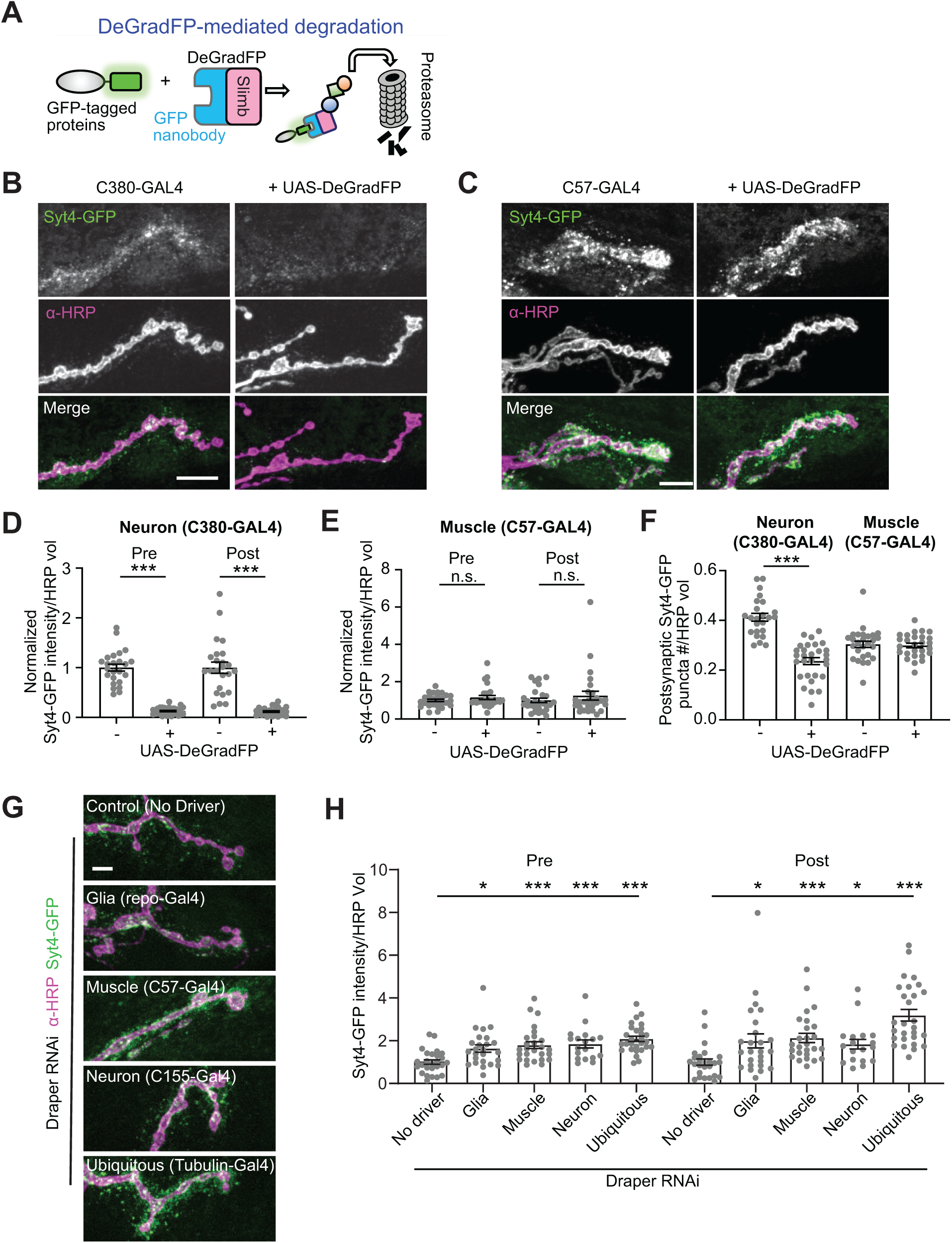
Neuronally derived EV cargoes are targeted for phagocytosis and are not detectable in the cytoplasm of recipient cells. **(A)** Schematic for DeGradFP system. **(B-C)** Representative images of Syt4-GFP with neuronal (C380-GAL4, **B**) or muscle (C57-GAL4, **C**) expressed DeGradFP. **(D)** Quantification of Syt4-GFP intensity from **(B)**. **(E)** Quantification of Syt4-GFP intensity from **(C)**. **(F)** Quantification of normalized presynaptic puncta number from **(B)** and **(C)**. **(G)** Representative confocal images of Syt4-GFP at muscle 4 NMJs following knockdown of Draper in different cell types. Outlines represent the neuronal membrane as marked from an HRP mask **(H)** Quantification of Syt4 puncta intensity. All scale bars = 10 µm. Intensity measurements are normalized to their respective controls. Data are represented as mean +/- s.e.m.; n represents NMJs. **p<0.01, ***p<0.001. **See Tables S1 and S3 for detailed genotypes and statistical analyses.**

Conversely, if EVs serve primarily as a proteostatic mechanism to shed neuronal Syt4 for subsequent uptake and degradation in recipient cells, then Syt4 would not need to be exposed to the muscle cytoplasm, as it could be taken up by phagocytosis in double membrane compartments for degradation via fusion with recipient cell lysosomes. Indeed, α- HRP positive neuronal “debris” is taken up via the phagocytic receptor Draper (Fuentes-Medel et al., 2009). This debris was not previously directly linked to EVs, though it co-localizes strongly with EV cargo (Walsh et al., 2021). We used cell type-specific Draper RNAi to test directly if known EV cargoes are cleared by Draper-dependent phagocytosis. First, we established that Draper is expressed in neurons, glia, and muscles at the NMJ. RNAi of Draper in each of these tissues depleted a subset of Draper immunostaining at the NMJ and axon, indicating that Draper is expressed in each of these cell types (**Fig. S5B-C**). We then quantified Syt4 levels in Draper RNAi larvae. Depletion of Draper in either muscles, glia, or neurons led to an increase in postsynaptic Syt4-GFP, indicating that the normal destination of Syt4 following release from the neuron is phagocytosis by multiple adjacent cell types (**Fig. 7G-H**). Interestingly, we found that EV cargoes also accumulated presynaptically upon Draper knockdown in glia, neurons, or muscles – this could be due to presynaptic reuptake of EVs by bulk endocytosis when they cannot be cleared by phagocytosis. Overall, these results show that Syt4 EVs are cleared by phagocytosis, similar to previously characterized α-HRP-labeled debris, but are not transported at detectable levels into the cytoplasm of muscle cells.

## Discussion

Here we show that the ESCRT pathway is required for EV cargo packaging at the *Drosophila* larval NMJ, and that these EVs are likely MVE-derived exosomes. We found that ESCRT depletion caused presynaptic accumulation of cargoes, defects in their axonal transport, and a dramatic loss of trans-synaptic transfer in EVs. Surprisingly, we found that this trans-synaptic transfer is not required for several physiological functions of EV cargoes Evi and Syt4. Further, neuronally-derived Syt4 is taken up by phagocytosis and could not be detected in the muscle cytoplasm, consistent with findings from HeLa cells that the majority of EV cargoes remain in the endosomal system of the recipient cell (O’Brien et al., 2022). Our results suggest that neuronal EV release for these cargoes at this developmental stage serves primarily proteostatic and not signaling functions.

### Functions of ESCRT in MVE biogenesis and EV release at synapses

We found that ESCRT is required for EV generation and release at the *Drosophila* larval NMJ. ESCRT components are also required for EV/exosome cargo release from primary neurons in culture (Gong et al., 2016) and Purkinje neurons *in vivo* (Coulter et al., 2018), but not for EV release of pathogenic APP variants or Evi from cell lines (Beckett et al., 2013; Cone et al., 2020), underscoring the importance of studying membrane traffic in *bona fide* neurons. Further, we found that upon ESCRT depletion, cargoes accumulate in intracellular compartments, suggesting that this population of NMJ EVs are MVE-derived exosomes rather than plasma membrane-derived microvesicles. This is consistent with the requirement for endosomal sorting machinery, such as retromer, in their regulation (Walsh et al., 2021).

One major open question is whether EV-precursor MVEs are generated on-demand in response to local cues at presynaptic terminals, or if they arise in response to global cues and are transported to synapses from other regions of the neuron. Answering this question will require tools to visualize the timecourse of MVE biogenesis in neurons, as have been developed in cultured non-neuronal cells (Wenzel et al., 2018). In addition, future methods (e.g. optogenetic) for acute and localized inhibition of ESCRT will reveal whether arrested structures first appear locally at the synapse and are only later transported into axons and cell bodies, and/or if they are generated far from the site of release at the synapse. These experiments will be critical for understanding when and where local or global signaling events impinge on EV biogenesis. Interestingly, activity-dependent delivery of Hrs to presynaptic terminals is critical for proteostasis of synaptic vesicle proteins (Birdsall et al., 2022; Sheehan et al., 2016). If MVEs are generated on-demand at synapses, Hrs transport could similarly underlie the activity-dependence of EV release, which has been reported in many (but not all) neuronal experimental systems, and remains poorly understood (Ataman et al., 2008; Faure et al., 2006; Lachenal et al., 2011; Lee et al., 2018; Vilcaes et al., 2021).

### Other synaptic functions of ESCRT

ESCRT is best-known for its functions in MVE biogenesis, but has many other potential synaptic roles including in autophagy, lipid transfer and membrane repair (Vietri et al., 2020). Tsg101 is involved in lipid transfer to mitochondria (Wang et al., 2021), but we did not detect obvious defects in mitochondria in motor neuron axons, as were seen in *Tsg101*-mutant *Drosophila* adult wing sensory neurons (Lin et al., 2021). Our results also show that the function of ESCRT in EV release is likely separate from its roles in autophagy, since several canonical autophagy mutants do not phenocopy presynaptic trapping of EV cargoes as seen upon ESCRT depletion, and *Hrs* mutants exhibit EV but not autophagic flux defects. Interestingly, we found that *atg* mutants led to a moderate presynaptic and postsynaptic reduction in levels of the EV cargo Nrg. This raises the possibility that other degradative pathways are upregulated at synapses when autophagy is blocked. Thus, while ESCRT has many cellular activities, our experiments separate these functions, and specifically narrow down its role in neuronal EV release.

Many organelles are selectively targeted for macroautophagy via compartment-specific receptors (Lamark and Johansen, 2021), but such a process has not been specifically described for neuronal endosomes/MVEs. Our data suggest that synapses use a proteostatic mechanism called endosomophagy or simaphagy that has been previously observed in cell culture (Migliano et al., 2023; Millarte et al., 2022; Wang et al., 2022; Zellner et al., 2021), adding to the numerous intersections between endolysosomal traffic and autophagy in neurons (Boecker and Holzbaur, 2019). We found that autophagy is induced in ESCRT mutant synapses, presumably to dispose of aberrant endosomes, with different outcomes in Tsg101^KD^ versus *Hrs* mutants: Tsg101^KD^ led to aberrant autophagic vacuoles and reduced autophagic flux, perhaps due to a secondary role for ESCRT-1/Tsg101 in phagophore closure or another step of autophagy (Takahashi et al., 2018). By contrast we found that *Hrs* mutants do not show these structures, either by light microscopy of the autophagic flux reporter GFP-mCherry-ATG8, or in previously published TEM of the NMJ (Lloyd et al., 2002). Instead, *Hrs* mutants exhibit induction of autophagy but normal autophagic flux in motor neurons, together with a moderate reduction in EV cargo levels. Together, these results suggest that aberrant EV-cargo-containing MVEs may be removed from *Hrs* mutant synapses by a compensatory, Tsg101-dependent autophagy pathway.

### Implications for the signaling roles of EVs

The majority of functional studies of EVs involve isolating EV subpopulations (at various degrees of homogeneity) from cell culture supernatants, applying them to target cells or tissues, and assessing their biological effects (Welsh et al., 2024). Additional mechanistic insight has been obtained by eliminating specific cargo molecules from the donor cells before EV isolation, to determine if these molecules are required for EV bioactivity. While these approaches are very useful for determining therapeutic uses for EVs, they have several major limitations for understanding their normal functions *in vivo*. First, it is difficult to determine the concentration of EVs that a target cell would normally encounter, in order to design a physiologically relevant experiment. Second, while these types of experiments inform what EVs can do, they do not show that EV transfer is necessary for that signaling function *in vivo*. Removing the signaling cargo from the donor cell also does not show the necessity of EV transfer for biological functions, since the cargo could be acting cell autonomously in the donor cell, or could signal to a neighboring cell by another trafficking route. Indeed, previous studies at the *Drosophila* larval NMJ, which has been an important model system for the *in vivo* functions of EV traffic, have conducted tests for EV cargo activity in *evi* or *rab11* mutants, though we and others have shown that this results in depletion of cargo from the presynaptic donor cell in addition to loss of EVs (Ashley et al., 2018; Blanchette et al., 2022; Koles et al., 2012; Korkut et al., 2009; Korkut et al., 2013; Walsh et al., 2021). Ultimately, determining if transfer of a cargo in EVs is necessary for its signaling function requires blocking EV transfer specifically, which we were able to achieve at ESCRT-depleted synapses.

Neuronally derived Wg is required and sufficient for synaptic growth, and functions together with glia-derived Wg to organize postsynaptic glutamate receptor fields (Kerr et al., 2014; Korkut et al., 2009; Miech et al., 2008; Packard et al., 2002). Therefore, if transsynaptic transfer of Evi and Wg in EVs was required for Wg signaling, we would expect to see a reduction in synaptic growth at ESCRT-depleted synapses, as well as disruptions in postsynaptic development and organization. However, we observed no significant change in bouton or active zone number relative to controls upon either Tsg101 or Hrs depletion, indicating that they do not phenocopy either *evi* or *wg* mutants, and that at least some Wg activity is maintained even when Evi-GFP transfer is strongly inhibited. *wg*-phenocopying defects in subsynaptic reticulum were also not observed by electron microscopy in CHMPIIB^intron5^ (West et al., 2015) or *Hrs*-mutant synapses (Lloyd et al., 2002), or in our data from ESCRT mutants. Similarly, Hsp90 mutants attenuate Evi EV release by disrupting MVE-plasma membrane fusion, but do not result in disruption of active zone or subsynaptic reticulum structure (Lauwers et al., 2018). Therefore, loss of EVs does not phenocopy many *wg* or *evi* defects, suggesting that the primary function of Evi is likely to traffic Wg to the presynaptic terminal and maintain its levels there, rather than specifically to mediate its release via EVs.

Importantly, it is likely that Hsp90 and ESCRT mutant synapses do secrete Wg, albeit by a non-EV mechanism (Beckett et al., 2013; Won and Cho, 2021). Interestingly, we found that Tsg101^KD^ leads to loss of Fz2-C nuclear import and an increase in baseline ghost boutons, consistent with some defects in Wg signaling. Our finding that Tsg101^KD^ causes additional membrane trafficking defects (e.g. autophagy) compared to *Hrs* suggests that non-EV modes of Wg release may be disrupted in this mutant. Likely possibilities include conventional secretion or secretory autophagy (Beckett et al., 2013; Chang et al., 2024; Won and Cho, 2021). However, since *Hrs* mutants do not show any deficit in Fz2-C nuclear import or ghost boutons despite exhibiting a similar loss of postsynaptically transferred Evi-GFP to Tsg101^KD^, we conclude that Wg signaling and EV release are separable functions.

Syt4 protein is thought to act in the postsynaptic muscle (Adolfsen et al., 2004; Barber et al., 2009; Harris et al., 2016), but its endogenous transcript is not expressed in this tissue, leading to the prevailing model of transynaptic transfer from the presynaptic neuron in EVs (Korkut et al., 2013). Our results show that transynaptic transfer in EV can be blocked without affecting the signaling activities of Syt4, and that the majority of postsynaptic Syt4 is not exposed to the muscle cytoplasm. The main evidence for a muscle requirement for Syt4 is that re-expression of Syt4 using muscle-specific GAL4 drivers is sufficient to rescue structural and functional plasticity defects of the *syt4* null mutant (Korkut et al., 2013; Piccioli and Littleton, 2014; Yoshihara et al., 2005). This is difficult to reconcile with our findings that Tsg101^KD^ and *Hrs* animals lack detectable postsynaptic Syt4, but do not phenocopy *syt4* mutants. There are several possible explanations for this conundrum. First, we cannot completely rule out the possibility that small amounts of residual Syt4 EVs are sufficient to drive a transynaptic signal. This is unlikely, since *nwk* and *rab11* mutants also have trace amounts of Syt4 postsynaptically, and do strongly phenocopy the *syt4* null mutant (presumably since they also deplete Syt4 from the presynaptic compartment) (Blanchette et al., 2022; Korkut et al., 2013; Walsh et al., 2021). Therefore, trace Syt4 is insufficient for signaling. Second, Syt4 could be transferred by a non-EV pathway such as conventional secretion, tunneling nanotubes, or cytonemes, and be distributed diffusely in the muscle such that we cannot detect its presence or degradation by cytoplasmic deGradFP (Dagar and Subramaniam, 2023; Daly et al., 2022)). Third, it is possible that the muscle GAL4 drivers and UAS lines used in these previous rescue studies have some leaky expression in the neuron. Fourth, ectopically muscle-expressed Syt4 might have a neomorphic function in the muscle that bypasses the loss of neuronal Syt4, or else it could be retrogradely transported to the neuron. Indeed, muscle-expressed Syt4 is localized in close apposition to the presynaptic membrane (Harris et al., 2016).

## Conclusions

Why are Evi and Syt4 trafficked into EVs, if not for a signaling function? Local EV release could serve as a proteostatic mechanism for synapse-specific control of signaling cargo levels, in cooperation with other degradative mechanisms. Our data show that EVs are taken up by glial, muscle, and neuronal phagocytosis, and that cargoes are protected from the muscle cytoplasm. Indeed, the amount of cargo loaded into EVs could be tuned by regulating endosomal sorting via retromer (Walsh et al., 2021), or by controlling the rate of endocytic flux into the Rab11-dependent recycling pathway (Blanchette et al., 2022). Our results also show that EVs are one of several intersecting and complementary mechanisms for synaptic proteostasis of membrane-bound cargoes; when EV release is reduced, we found that compensatory autophagy pathways are upregulated to degrade unwanted endosomal components. Further, endosomes that are not competent for EV biogenesis can be targeted for dynein-mediated retrograde transport (Blanchette et al., 2022), perhaps to bring cargoes to the cell body where lysosomal degradation is more active (Ferguson, 2018). Through these mechanisms, neurons might achieve local control of synaptogenic or plasticity-inducing signaling pathways, in a much more rapid and spatially controlled fashion than transcriptional or translational regulation.

Importantly, our results do not rule out signaling functions for Syt4 or Evi/Wg EVs in other contexts or neuronal cell types, or for other EV cargoes. For example, ESCRT disruption suppresses the pathological functions of APP in *Drosophila*, perhaps due to its reduced propagation in EVs (Zhuang et al., 2023). Indeed, extensive evidence supports signaling and pathological functions for neuronal EVs in multiple contexts (Gassama and Favereaux, 2021; Lizarraga-Valderrama and Sheridan, 2021; Schnatz et al., 2021). However, our data warrant future hypothesis-challenging experiments for EV functions using membrane trafficking mutants that disrupt EV release specifically.

## Materials and Methods

### Drosophila culture

Flies were cultured using standard media and techniques, except larvae for **Fig. 5F-G** (FzC-2 nuclear import), which were cultured on Gerber Natural for Baby, Peach, 2^nd^ Foods (Sitter). Flies used for experiments were maintained at 25°C, except for experiments using Shrub-RNAi, which were maintained at 20°C. Suitable reagents were not available to assess the extent of Tsg101 or Shrub knockdown. However, given that we observe very similar phenotypes for ESCRT RNAi, genomic mutants, and dominant negative mutants (**Figs. 1-2**), we conclude that these RNAi tools phenocopy strong loss-of-function, and that the phenotypes we observe are specific. For detailed information on fly stocks used, see **Table S1**, and for detailed genotype information for each figure panel, see **Table S3**.

### Immunohistochemistry

Wandering 3^rd^ instar larvae were dissected in HL3.1 and fixed in HL3.1 with 4% paraformaldehyde for 45 minutes (except **Figs. 2A,B,D,E, Fig. 7** and **Fig. S5** which were fixed for 10 minutes). For α-Fz2 staining, wandering 3^rd^ instar larvae were dissected in 0 mM Ca^2+^ modified *Drosophila* saline (Restrepo et al., 2022) and fixed in 4% paraformaldehyde for 20 minutes. Washes and antibody dilutions were conducted using PBS containing 0.2% Triton X-100 (0.2% PBX), except Fz2-C stain washes and antibody dilutions which were conducted using PBS containing 0.3% Triton X-100. Primary antibody incubations were conducted overnight at 4°C, and secondary antibody incubations for 1-2 hours at room temperature. α-HRP incubations were conducted either overnight at 4°C or for 1-2 hours at room temperature. Prior to imaging, fillets were mounted on slides with Vectashield (Vector Labs) or Abberior Liquid Antifade (Abberior). For detailed information on antibodies used in this study, see **Table S2**.

### Electron microscopy

Wandering 3^rd^ instar larvae were dissected and fixed in 1% glutaraldehyde and 4% paraformaldehyde in 1% (0.1M) sodium cacodylate buffer overnight at 4°C. Samples were postfixed in 1% osmium tetroxide and 1.5% potassium ferrocyanide for 1 h, then 1% aqueous uranyl acetate for 0.5 h. Stepwise dehydration was conducted for 10 min each in 30%, 50%, 70%, 85%, and 95% ethanol, followed by 2× 10 min in 100% ethanol. Samples were transferred to 100% propylene oxide for 1 h, then 3:1 propylene oxide and 812 TAAB Epon Resin (epon, TAAB Laboratories Equipment Ltd.) for 1 h, then 1:1 propylene oxide:epon for 1 h and then left overnight in a 1:3 mixture of propylene oxide:epon. Samples were then transferred to fresh epon for 2 h. Samples were then flat-embedded and polymerized at 60°C for 48 h, and remounted for sectioning. 70-µm-thin sections were cut on a Leica UC6 Ultramicrotome (Leica Microsystems), collected onto 2X1 mm slot grids coated with formvar and carbon, and then poststained with lead citrate (Venable and Coggeshall, 1965). Grids were imaged using a FEI Morgagni transmission electron microscope (FEI) operating at 80 kV and equipped with an AMT Nanosprint5 camera.

### Activity-induced synaptic growth

High K^+^ spaced stimulation was performed as described (Piccioli and Littleton, 2014). Briefly, 3^rd^ instar larvae were dissected in HL3 saline (Stewart et al., 1994) at room temperature (in mM, 70 NaCl_2_, 5 KCl, 0.2 CaCl_2_, 20 MgCl_2_, 10 NaHCO_3_, 5 trehalose, 115 sucrose, and 5 HEPES (pH=7.2)). Dissecting pins were then moved inward to relax the fillet to 60% of its original size, and then stimulated 3 times in high K^+^ solution (in mM, 40 NaCl_2_, 90 KCl, 1.5 CaCl_2_, 20 MgCl_2_, 10 NaHCO_3_, 5 trehalose, 5 sucrose, and 5 HEPES (pH=7.2)) for 2 minutes each, with 10-minute HL3 incubations in between stimulation while on a shaker at room temperature. Following the 3^rd^ and final stimulation, larvae were incubated in HL3 (approximately 2 minutes) and stretched to their initial length. Mock stimulations were performed identically to the high K^+^ stimulation assay, except HL3 solution was used in place of high K^+^ solution. Larvae were then fixed in 4% PFA in HL3 solution for 15 minutes and then stained and mounted as above.

### Electrophysiology

Wandering 3^rd^ instar larvae were dissected in HL3 saline. Recordings were taken using an AxoClamp 2B amplifier (Axon Instruments, Burlingame, CA). A recording electrode was filled with 3M KCl and inserted into muscle 6 at abdominal segments A3 or A4. A stimulating electrode filled with saline was used to stimulate the severed segmental nerve using an isolated pulse stimulator (2100; A-M Systems). HFMR was induced by four trains of 100 Hz stimuli spaced 2 s apart in 0.3 mM extracellular Ca^2+^. Miniature excitatory junctional potentials (minis) were recorded 2 min before and 10 min after HFMR induction for Tsg101. Many *hrs* mutant larvae did not maintain quality mini recordings over 10 minutes, so we recorded for only 5 minutes. Mini frequency at indicated time points was calculated in 10-s bins. Fold enhancement was calculated by normalizing to the baseline mini frequency recorded prior to HFMR induction. Analyses were performed using Clampfit 10.0 software (Molecular Devices, Sunnyvale, CA). Each n value represents a single muscle recording, with data generated from at least six individual larvae of each genotype arising from at least two independent crosses. Resting membrane potentials were between -50 mV and -75 mV and were not different between genotypes. Input resistances were between 5 MΩ and 10 MΩ and were not different between genotypes.

### Imaging and quantification

*Acquisition:* Analysis of EV cargoes and bouton morphology were conducted from larval abdominal muscles and segments as indicated in **Table S3**. Z-stacks were acquired using a Nikon Ni-E upright microscope equipped with a Yokogawa CSU-W1 spinning disk head, an Andor iXon 897U EMCCD camera, and Nikon Elements AR software. A 60X (n.a. 1.4) oil immersion objective was used to image NMJs, cell bodies, and fixed axons. Data in **Fig. S4C-D** were acquired with a Zeiss LSM 800 confocal microscope using a 40x (n.a. 1.4) oil immersion objective and Zen Black 2.3 software. For colocalization and puncta analysis branches from muscle 6/7 NMJ from segments A2 and A3 were imaged using Zen Blue software on a Zeiss LSM880 Fast Airyscan microscope in super resolution acquisition mode, using a 63X (n.a. 1.4) oil immersion objective.

Data in **Fig. S1B-E** were acquired on an Abberior FACILITY line STED microscope with 60x (NA1.3) silicone immersion objective, pulsed excitation lasers (561 nm and 640 nm), and a pulsed depletion laser (775 nm) to deplete all signals. Nrg was labeled with STAR ORANGE (Abberior, Inc.) and APP-GFP was labeled with anti-GFP antibodies and STAR RED-labeled secondaries (Abberior, Inc.). Pixel size was set to 40 nm, and single 2D slices were acquired of terminal branches of NMJs that lay in a single focal plane. α-HRP signal was detected using conventional confocal imaging.

For axonal transport, wandering 3^rd^ instar larvae were dissected one at a time in HL3.1. For APP-GFP, larvae were mounted between a slide and coverslip in HL3.1. For Mito-GFP axons the larvae were pinned in a sylgard coated dish covered with HL3.1. Dissection and imaging for each larva was completed within 30 minutes. Timelapse images were taken on the same Nikon Ni-E microscope described above. Images were taken of axon bundles proximal to the ventral ganglion (roughly within 100-300 µm). For APP transport, timelapse images were acquired for 3 minutes using 60X (n.a. 1.4) oil immersion objective. For mitochondria timelapse, images were acquired for 7 minutes using 60X (n.a. 1.0) water immersion objective. 9 Z slices were collected per frame (Step size 0.3 µm, with no acquisition delay between timepoints, resulting in a frame rate of 2.34-2.37 sec/frame). To visualize moving particles for mitochondria, a third of the axon in the field of view was photobleached using an Andor Mosaic digital micromirror device operated by Andor IQ software, to eliminate fluorescence from stationary particles that would interfere with visualization of particles moving into the bleached region. Image acquisition settings were identical for all images in each independent experiment.

#### EV cargo quantification and colocalization

Volumetric analysis was performed using Volocity 6.0 software. For each image, both type 1s and 1b boutons were retained for analysis while axons were cropped out, (except in limited cases where 1s boutons were very faint and therefore were cropped as their inclusion would cause the HRP threshold to be excessively dilated for other branches). The presynaptic volume was defined by an HRP threshold, excluding objects smaller than 7 µm^3^ and closing holes. The postsynaptic region was defined by a 3.3 µm dilation of the HRP mask. However, for Evi-GFP, where the presynaptic signal vastly exceeded postsynaptic signal, we analyzed only the distal 2.9 µm of this postsynaptic dilation region to eliminate the bleed-over haze from the presynaptic signal. EV cargo and Rab signals were manually thresholded to select particles brighter than the muscle background. EV cargo integrated density in these thresholded puncta was normalized to the overall presynaptic volume. These values were further normalized to the mean of the control to produce a “normalized puncta intensity” value for each NMJ. For colocalization, the overlap of the two channels was measured in Volocity 6.0 and used for calculation of Mander’s coefficients.

To perform particle-based analyses of EV puncta density and width, 2D-STED micrographs were denoised using Noise2Void (Krull et al., 2019). Briefly, a model was trained using the Nrg channel, using 10 control and 10 Tsg101^KD^ images as a training set. This model was used to denoise both Nrg and APP channels. Presynaptic regions were segmented using a conventional confocal image of HRP, as described above. This mask was dilated by 3 µm to generate a postsynaptic mask containing the vast majority of EV signal, from which a 10% dilation of the presynaptic mask was subtracted to remove any presynaptic signal. Finally, the postsynaptic mask was further dilated by 10% to generate a 300 nm buffer, and the remainder of the image was defined as background (e.g. nonspecific antibody signal) (BG, see **Fig. S1C** for schematic). To detect particles, each channel was independently rescaled from 0-1 (min and max pixel values), processed by a Mexican Hat filter (radius=4), and local intensity maxima were detected using a prominence value of 1.25. Local maxima were used as seeds to fit a 2D gaussian on the original (unscaled) denoised images, using the plugin GaussFit on Spot (https://imagej.net/ij/plugins/gauss-fit-spot/index.html). Parameters for fitting were as follows: shape=Circle fitmode=NelderMead rectangle=2.5 pixel=40 max=500 cpcf=1 base=0.

#### Quantification of electron micrographs

A single medial section of each bouton was selected for analysis. Two experimenters, blinded to genotype, together recorded the presence of autophagic vacuoles, including phagopores (double or dense membrane but not closed; note that depending on the plane of section, some of these may appear as autophagosomes), autophagosomes (contents with similar properties to the cytoplasm, fully enclosed in the section by a double membrane), and autolyososomes (contents are electron dense) (Lucocq and Hacker, 2013; Nagy et al., 2015). We also evaluated whether boutons lacked subsynaptic reticulum, or featured postsynaptic pockets (electron-lucent areas extending at least 300 nm from the presynaptic membrane (Packard et al., 2002).

#### Quantification of GFP-mCherry-Atg8 distribution

A single field-of-view confocal stack (62×62 µm) from the larval ventral ganglion, containing 10-15 Vglut-expressing cell bodies, was manually thresholded in Volocity 6.0 software to segment and measure the volume and integrated fluorescence density of soma, GFP puncta, and mCherry puncta. The overlap between the GFP and mCherry channels was used for the calculation of the Mander’s coefficient (fraction of total mCherry-puncta integrated density found in the GFP-puncta positive volume).

#### Axon and cell body measurements

To measure intensity of EV cargoes, axons proximal to the ventral ganglion (within 100-300 µm) were imaged as described above. Images were analyzed in Fiji by making sum projections, cropping out unwanted debris or other tissue and generating a mask from the α-HRP signal. The total intensity of the EV cargo was then measured within the masked HRP area. For cell bodies, EV cargo intensity was measured from a middle slice through the motor neuron cell body layer of the ventral ganglion using Fiji.

#### Quantification of live axonal trafficking of APP-GFP and Mito-GFP puncta

To quantify APP-GFP and mitochondria dynamics in live axons, maximum intensity projections of time course images were processed in Fiji to subtract background and adjust for XY drift using the StackReg plugin. Kymographs were generated from 1-4 axons per animal using the Fiji plugin KymographBuilder. Kymographs were blinded and number of tracks were manually counted. The minimum track length measured was 3 μm with most tracks above 5 μm. Velocity was measured by calculating the slope of the identified tracks.

#### Bouton quantification

The experimenter was blinded to genotypes and then manually counted the total number of type 1 synaptic boutons on the NMJ on muscle 6 and 7 in the abdominal segments A2 and A3 of third instar wandering larvae. A synaptic bouton was considered a spherical varicosity, defined by the presence of the synaptic vesicle marker Synaptotagmin 1, the active zone marker Bruchpilot (Brp) or the neuronal membrane marker Hrp. For quantifying ghost boutons (basal and activity-induced), the experimenter was blinded to genotype and condition and ghost boutons were quantified as α-HRP-positive structures with a visible connection to the main NMJ arbor, and without α-DLG staining. For quantifying DLG “featheriness”, the experimenter was blinded to genotype and scored the number of NMJs with at least one region of fenestrated Dlg that extended far from the bouton periphery.

#### Active zone quantification

To count the active zones in fluorescence micrographs, Brp-stained punctae were assessed on maximum intensity projection images. The Trainable Weka Segmentation (TWS) machine-learning tool (https://doi.org/10.1093/bioinformatics/btx180) in Fiji was used to manually annotate Brp-positive punctae with different fluorescence intensities, and to train a classifier that automatically segmented the Brp-positive active zones. The objects segmented via the applied classifier were subjected to Huang auto thresholding to obtain binary masks. Next, we applied a Watershed processing on the binary image, to improve the isolation of individual neighboring active zones from the diffraction limited images. We performed particle analysis on the segmented active zones and obtained their number, area, and integrated density. To determine the NMJ area using TWS, we trained the classifier by annotating the HRP positive NMJ on maximum intensity projections of the HRP channel. Axons were manually cropped from the image before TWS. The segmented HRP area was subjected to Huang auto thresholding, the binary masks were selected and the NMJ area was obtained via the “Analyze particle” function in FIJI of particles larger than 5 µm (to eliminate from the analysis residual HRP EV debris segmented in a very few images).

#### Quantification of Frizzled2 C-terminus nuclear localization

To quantify Fz2-C puncta, muscles 6 and 7 were imaged from segments A2 and A3 in larvae where the experimenter was blinded to genotype. Muscle nuclei were identified by the boundaries of LamC staining (which recognizes the nuclear envelope). Nuclear puncta were quantified as aggregates of staining that exceeded the size and fluorescence intensity of non-specific background staining from the rabbit α-Fz2-C antibody. No nuclei were excluded from quantification and all nuclei pooled for final statistical analysis. In all genotypes, >250 individual nuclei were scored.

#### Quantification of Draper knockdown

To analyze Draper levels in axonal bundles (which include neurons and glia), a region of axon proximal to muscle 4 was cropped to 100×100 pixels in a 3D spinning disk confocal Z-stack, using Fiji. This image was masked on the HRP channel and the mean α-Draper intensity was calculated in the 3D volume. To analyze Draper levels at the NMJ (which includes the presynaptic neuron and adjacent or optically overlapping postsynaptic muscle membrane), images were cropped to include only type 1b bouton branches (excluding axon bundles, axon, type 1s bouton branches, or any non-bouton material). The image was masked on the HRP channel, dilated by 0.22 µm, and the mean α-Draper intensity was calculated in the 3D volume.

### Statistics

All statistical measurements were performed in GraphPad Prism (see **Table S3**). Comparisons were made separately for presynaptic and postsynaptic datasets, due to differences between these compartments for intensity, signal-to-noise ratio, and variance. Datasets were tested for normality, and statistical significance was tested as noted for each experiment in **Table S3**. Statistical significance is indicated as *P < 0.05; **P < 0.01; ***P < 0.001.

## Acknowledgements

We thank the Developmental Studies Hybridoma Bank created by the NICHD of the NIH, and the Bloomington *Drosophila* Stock Center (Indiana University, Bloomington, IN, NIH P40OD018537). This work was supported by grant 1S10OD034223-01 for the Abberior Facility Line STED microscope, housed in the Brandeis Light Microscopy Core Facility. We thank Berith Isaac and the Brandeis Electron Microscopy Facility for assistance with EM, Michael Boutros, Josie Clowney, and Tor Erik Ruston for fly lines, and Harald Stenmark for discussion of simaphagy. This work was supported by NINDS grants R01 NS103967 to A.A.R., F32 NS120909 to E.C.D., and R01 NS110907 to T.J.M.

## Author contributions

ECD, KPH, KK, MFP, CRB, BAS, TJM, and AAR designed the study and experiments. ECD, KPH, KK, MFP, BE, CRB, MR, RCS, TJM conducted the experiments. ECD, KPH, KK, MFP, BE, CRB, SJDS, MR, RCS, TJM and AAR performed the analyses. ECD and AAR wrote the manuscript, and all authors edited the manuscript. This article contains supporting information (5 Supplemental Figures and 3 tables).

---

**Figure S1:**
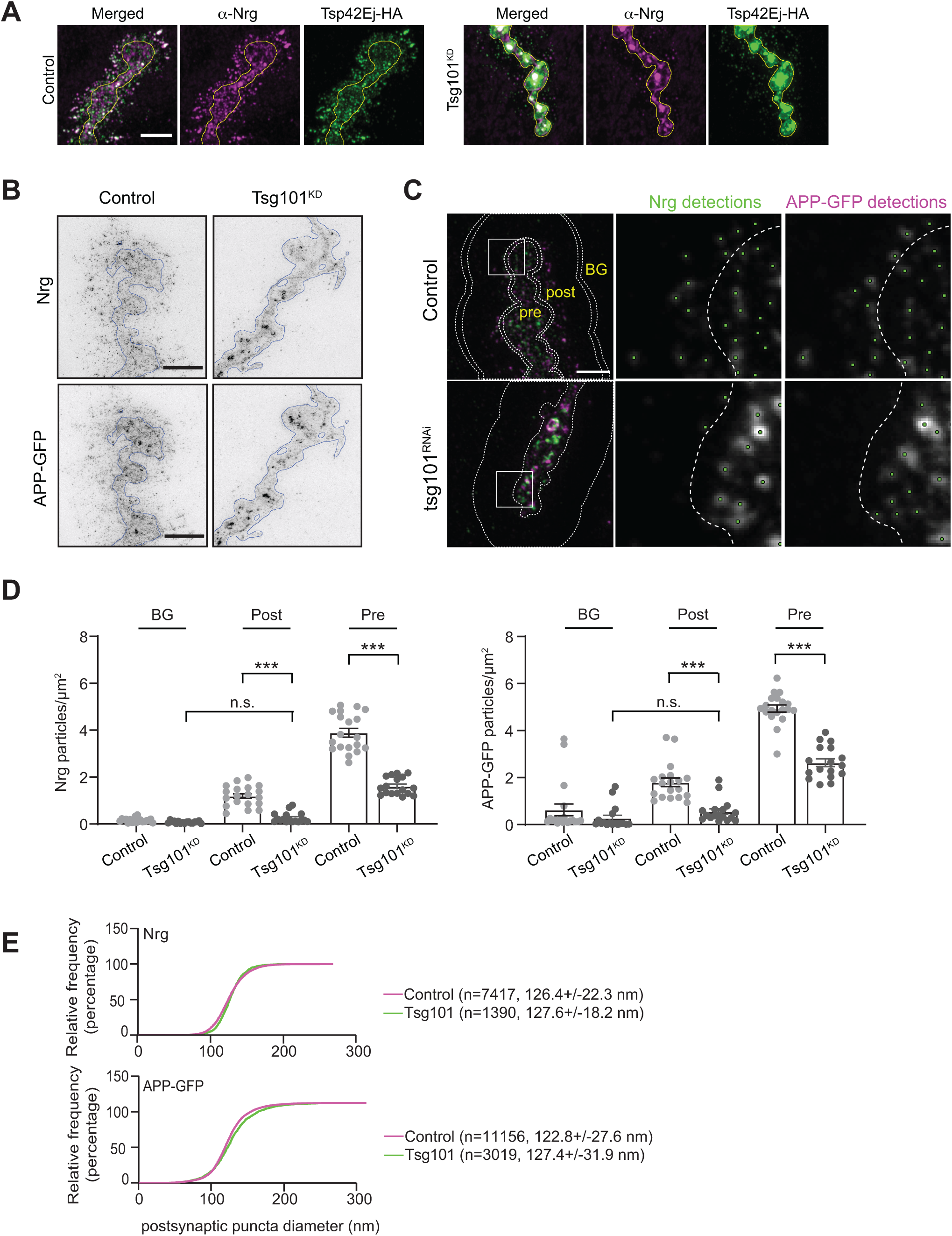
Characterization of EV structures upon Tsg101 depletion (associated with. **Figure 1). (A)** Representative Airyscan images of larvae expressing UAS-Tsp42Ej-HA, and labeled with α- HA and α-Nrg. Scale bar is 5 µm. Yellow outline represents the neuronal membrane as marked from an HRP mask. **(B)** Representative 2D-STED images of muscle 6/7 labeled with α-GFP and α-Nrg antibodies. Scale bar is 2.5 µm. **(C)** Noise2Void denoised images and depiction of image regions used for quantification (left panel, scale bar is 2.5 µm). Pre: presynaptic; Post: Postsynaptic; BG: Background. Buffers (between double lines in the top left panel) generated by a 10% dilation of the presynaptic or postsynaptic area were used to eliminate signal that overlapped between regions. Boxes indicate zoomed areas in middle and right panels showing automated spot detections (green dots) and the presynaptic boundary (dotted line). **(D**) Quantification of APP-GFP and Nrg puncta number. Data are represented as mean +/- s.e.m.; n represents NMJs; ***p<0.001. **(E)** Cumulative distribution of Nrg and APP puncta diameter. Graph shows fraction of particles under the indicated size; numbers indicate mean and standard deviation of all detected puncta. **See Tables S1 and S3 for detailed genotypes and statistical analyses.**

**Figure S2:**
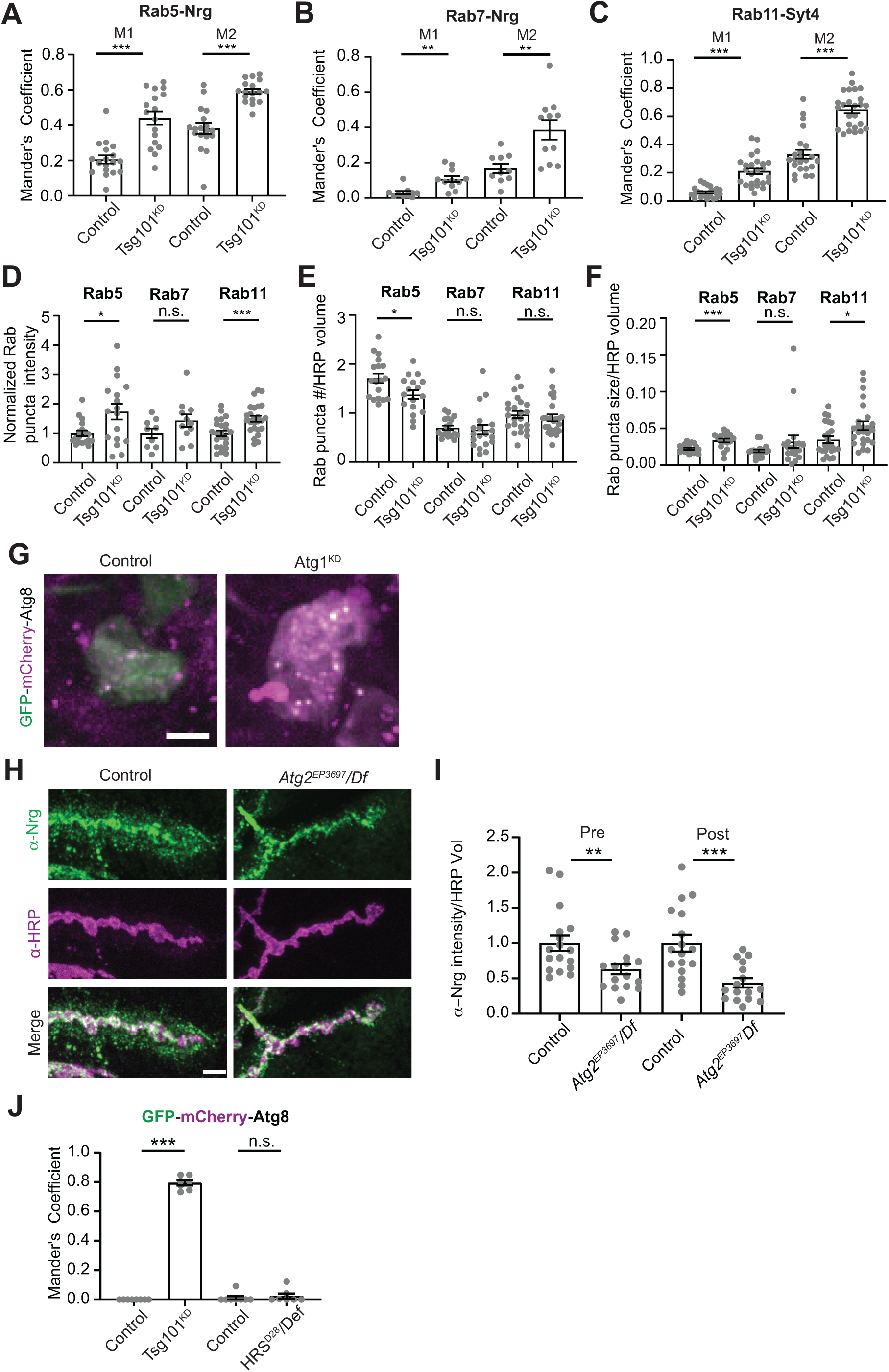
Quantification of endosomal accumulation and autophagy controls (associated with Figure 3). **(A-C)** Quantification of co-localization of Nrg or Syt4 and Rab GTPases upon neuronal Tsg101^KD^ (representative images in **Fig. 3A**). Mander’s coefficient for the colocalization of Nrg and Rab5 **(A)**, Nrg and Rab7 **(B)**, Syt4 and Rab11 **(C)**, where M1 indicates the fraction of EV cargo in the Rab-positive thresholded area and M2 is the fraction of the Rab marker in the EV cargo-positive thresholded area. **(D-F)** Quantification of Rab compartment properties: **(D)** normalized Rab puncta intensity, **(E)** density of Rab puncta in the presynaptic compartment, and **(F)** average size of Rab puncta. **(G)** Representative confocal images of motor neuron cell bodies to validate that pan-neuronal Atg1 RNAi effectively blocks autophagic flux, assessed by GFP-mCherry-Atg8. **(H)** Representative confocal images of Nrg in muscle 6/7 NMJs **(I)** Quantification of Nrg intensity from **(H). (J)** Colocalization of GFP and mCherry in cell bodies from motor neurons expressing GFP-mCherry-Atg8 (representative images in **Fig 3E**). All scale bars = 5 µm. Data are represented as mean +/- s.e.m.; n represents NMJs in **(A-F, I)** and animals in **(J**). Intensity measurements (**D, I**) are normalized to their respective controls. *p<0.05, **p<0.01, ***p<0.001. **See Tables S1 and S3 for detailed genotypes and statistical analyses.**

**Figure S3.**
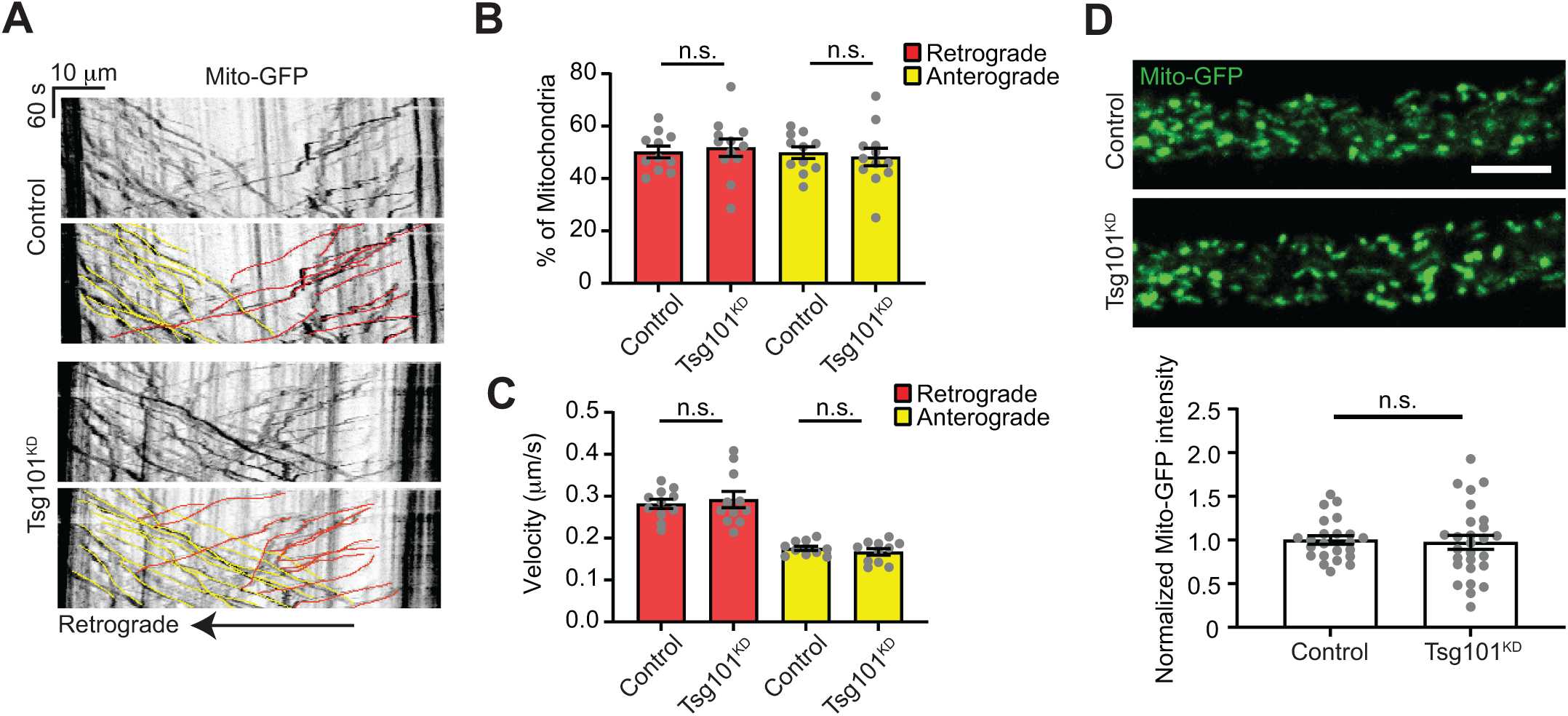
Controls for axonal transport in Tsg101^KD^ larvae (associated with Figure 4). **(A)** Representative kymographs showing tracks of Mito-GFP in axonal region proximal to the ventral ganglion, following photobleaching. Lower panels show color coded traces. **(B)** Percent of mitochondria tracks moving retrograde and anterograde. **(C)** Velocities of mitochondria tracks. **(D) (Top)** Representative images of the first frame of Mito-GFP videos. Scale bar = 10 µm. **(Bottom)** Quantification of Mito-GFP intensity. Data are represented as mean +/- s.e.m.; n represents axons. Intensity measurements are normalized to their respective controls. **See Tables S1 and S3 for detailed genotypes and statistical analyses.**

**Figure S4.**
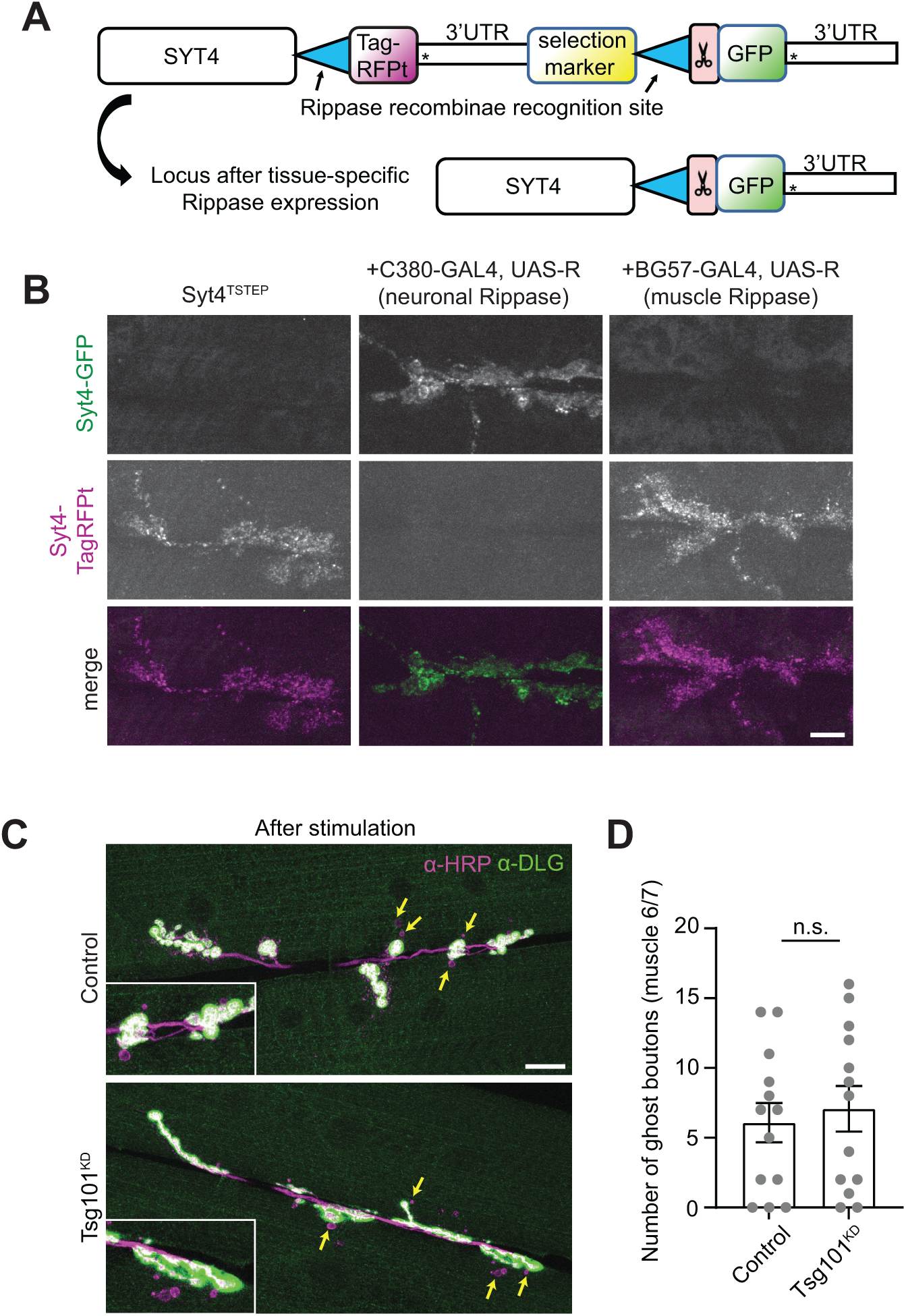
Additional controls showing presynaptic source of Syt4 and structural plasticity upon Tsg101^KD^ (associated with Figure 6). **(A-B)** Syt4 protein is derived from the presynaptic neuron. **(A)** Schematic for Tissue-Specific Tagging of Endogenous Proteins (T-STEP). Scissors indicate a Prescission protease cleavage site and * indicates stop codons. **(B)** Representative confocal images from muscle 6/7, showing Syt4^TSTEP^ expressed from its endogenous promoter, and switched from TagRFPt to GFP using either presynaptically (neuronal, C380-GAL4) or postsynaptically (muscle, C57-Gal4)-expressed recombinase (Rippase). Scale bar = 10 µm. **(C)** Representative confocal images from muscle 6/7 in spaced K^+^-stimulated larvae. Arrows indicate ghost boutons. Scale bar = 20 µm. **(D)** Quantification of ghost bouton numbers per NMJ. Scale bars = 10 µm. Data is represented as mean +/- s.e.m.; n represents NMJs. **See Tables S1 and S3 for detailed genotypes and statistical analyses.**

**Figure S5.**
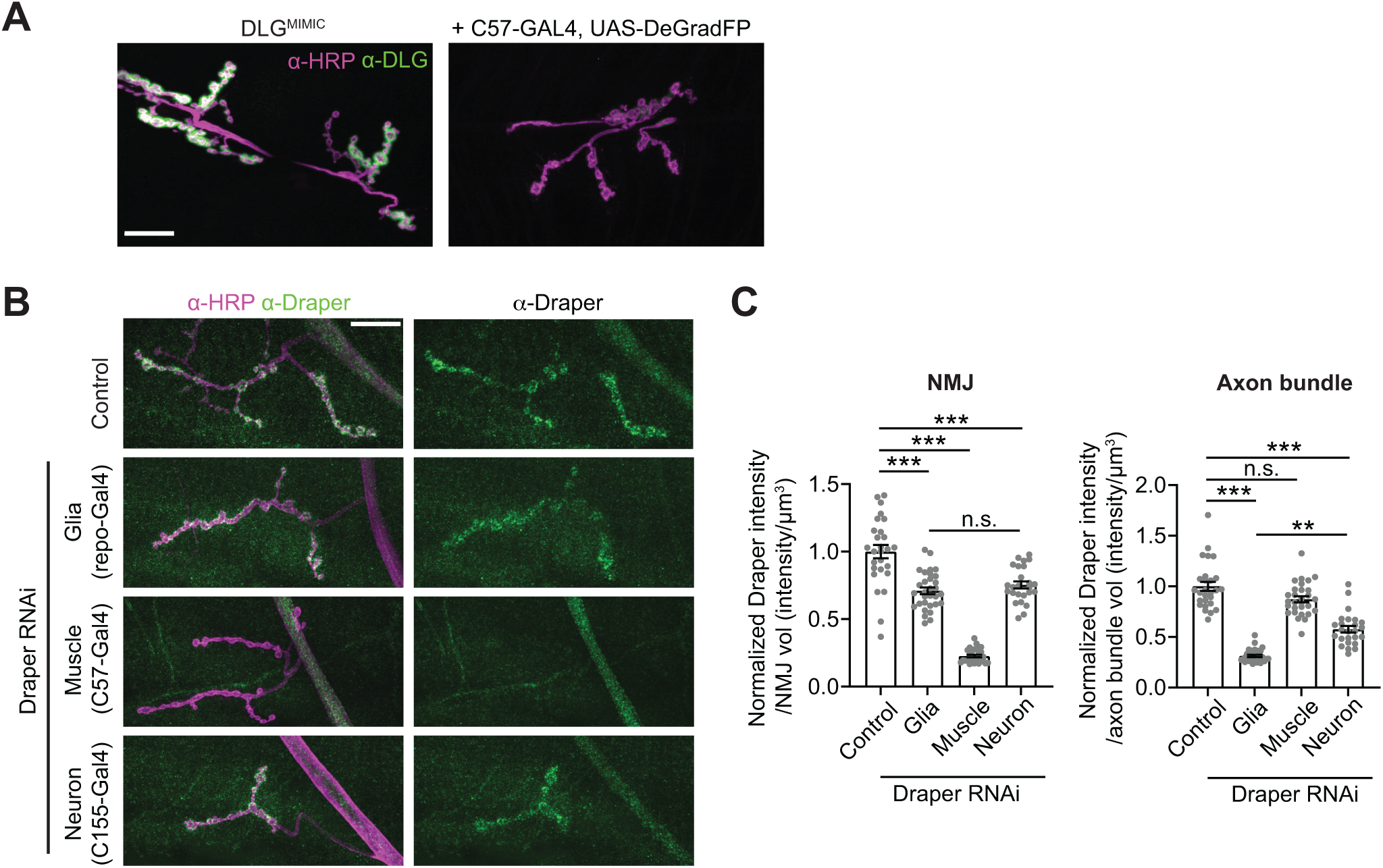
Controls for DeGradFP and validation of Draper RNAi (associated with Figure 7). **(A)** Representative images of Dlg^MiMIC^ (a postsynaptically-localized GFP knock-in) with muscle-expressed DeGradFP. **(B)** Representative confocal images of muscle 4 NMJs labeled with α-Draper antibodies**. (C)** Quantification of α-Draper intensity at NMJs and axon bundles proximal to the NMJ upon Draper RNAi under the control of the indicated drivers. Axon bundles represent a combination of glial and neuronal signal; NMJs represent a combination of neuronal and muscle signal. Scale bars are 20 µm. Intensity measurements are normalized to their respective controls. Data are represented as mean +/- s.e.m.; n represents NMJs. **p<0.01, ***p<0.001. **See Tables S1 and S3 for detailed genotypes and statistical analyses.**

**Table S1:** *Drosophila* mutants and genetic tools. Chromosome for transgene insertion indicated in roman numerals.

|  |  |  |
| --- | --- | --- |
| *GAL4<sup>C380</sup>* (X) | (Budnik et al., 1996) | Flybase ID:FBti0016294 |
| *GAL4<sup>C155</sup>* (X) | (Lin and Goodman, 1994) | Flybase ID: FBti0002575 |
| *GAL4<sup>Vglut</sup>* (X) | {Daniels, 2008 #424} | Flybase ID: FBti0129146 |
| *GAL4<sup>C57</sup>* (III) | (Budnik et al., 1996) | Flybase ID: FBti0016293 |
| *GAL4<sup>repo</sup>* (III) | (Sepp et al., 2001) | Flybase ID: FBti0018692 |
| *GAL4<sup>Tub</sup>* (III) | (Lee and Luo, 1999) | Flybase ID: FBal0097158 |
| *w<sup>1118</sup>* | (Hazelrigg et al., 1984) | Flybase ID: FBal0018186 |
| *Syt4<sup>EGFP-KI</sup>* | (Walsh et al., 2021) | Flybase ID: FBal0368282 |
| UAS-Evi-EGFP (II) | (Bartscherer et al., 2006) | Flybase ID: FBal0194740 |
| UAS-APP-EGFP (II) | (Walsh et al., 2021) | Flybase ID: FBal0368284 |
| pUAS-Tsp42Ej-3xHA (AttP1) | (Walsh et al., 2021) | Flybase ID: FBal0368283 |
| GFP-Rab5-KI (II) | (Fabrowski et al., 2013) | Flybase ID: FBal0288693 |
| YFP-myc-Rab7-KI (III) | (Dunst et al., 2015) | Flybase ID: FBal0314198 |
| UAS-Mito-HA-GFP (II) | (Rizzuto et al., 1995) | Flybase ID: FBti0040803 |
| UAS-GFP-mCherry-Atg8 (II) | (Nezis et al., 2010) | Flybase ID: FBti0147141 |
| UAS-mCherry-RNAi (III) | (Perkins et al., 2015) | Flybase ID: FBal0260847 |
| UAS-Luciferase-RNAi (III) | (Perkins et al., 2015) | Flybase ID: FBti0143388 |
| UAS-Tsg101-RNAi<sup>GLV21075</sup> (III) (Tsg101<sup>KD</sup>) | (Perkins et al., 2015) | Flybase ID: FBti0144671 |
| UAS-Shrub-RNAi<sup>HMS01767</sup> (II) (Shrub<sup>KD</sup>) | (Perkins et al., 2015) | Flybase ID: FBti0149514 |
| UAS-Vps4<sup>DN</sup> (II) | (Rodahl et al., 2009) | Flybase ID: FBal0221338 |
| *Hrs<sup>D28</sup>* | (Lloyd et al., 2002) | Flybase ID: FBal0039519 |
| *evi<sup>2</sup>* | (Bartscherer et al., 2006) | Flybase ID: FBal0194741 |
| *Syt4<sup>BA1</sup>* | (Adolfson et al., 2004) | Flybase ID: FBal0191284 |
| Df(2L)Exel6277 (removes Hrs) |  | Flybase ID: FBab0037889 |
| UAS-Atg1-RNAi<sup>GL00047</sup> (III) (Atg1<sup>KD</sup>) |  | Flybase ID: FBti0144152 |
| *Atg2<sup>EP3697</sup>* | (Shen and Ganetzky, 2009) | Flybase ID: FBti0011778 |
| Df(3L)Exel6091 (removes Atg2) |  | Flybase ID: FBab0038111 |
| UAS-DegradGFP (II) (nSLimb-vhhGFP4) | (Caussinus et al., 2011) | Flybase ID: FBal0299649 |
| Dlg1-MiMIC (X) | (Nagarkar-Jaiswal et al., 2015) | Flybase ID: FBti0168863 |
| UAS-Rippase (III) | (Nern et al., 2011) | Flybase ID: FBst0055795 |
| *Syt4<sup>TSTEP</sup>* | (Walsh et al., 2021) | Flybase ID: FBti0216043 |
| UAS-Draper-RNAi (II) | (MacDonald et al., 2006) | Flybase ID: FBal0258016 |

**Table S2:** Antibodies.

| REAGENT | SOURCE | IDENTIFIER | Concentration |
| --- | --- | --- | --- |
| **Antibodies** |  |  |  |
| α-HRP-647 | Jackson ImmunoResearch | 123-605-021 | 1:250-1:500 |
| α-Rab11 | BD Biosciences | BD 610657 | 1:100 |
| α-Dlg (mouse) | (Parnas et al., 2001), DSHB | 4F3 | 1:500 |
| α-Dlg (rabbit) | (Koh et al., 1999) | | 1:10000 |
| α-Nrg | (Hortsch et al., 1990), DSHB | BP104 | 1:100 |
| α-BRP | (Wagh et al., 2006), DSHB | nc82 | 1:100 |
| α-Draper | DSHB | 5D14 | 1:200 |
| α-HA | | HA.11 | 1:500 |
| α-GFP | Abcam | Ab6556 | 1:500 |
| α-Frizzled-C | (Mathew et al., 2005) | | 1:300 |
| α-rabbit STAR RED | Aberrior | 1002 | 1:250 |
| α-mouse STAR ORANGE | Aberrior | 1001 | 1:250 |

**Table S3:** Genotypes and Statistics by Dataset. Experiments were done at muscle 6/7 unless otherwise noted All larvae are male unless otherwise noted Analysis performed in Volocity unless otherwise noted Presynaptic volume: α-HRP objects > 7μm^3^ Postsynaptic volume: 3μm dilated from presynaptic volume A: Sum intensity of signal in thresholded objects in presynaptic volume, normalized to presynaptic volume B: Sum intensity of signal in thresholded objects in postsynaptic volume, normalized to presynaptic volume n.s.: not significantly different; SDCM: spinning disk confocal microscopy; LSCM: laser scanning confocal microscopy; TEM: transmission electron microscopy; STED: Stimulated Emission Depletion Microscopy

| Figure | Genotype/Conditions | N | Measurement | Statistical Test(s) |
| --- | --- | --- | --- | --- |
| **1A, E**<br>**SDCM** | Gal4<sup>C380</sup>/Y;; Syt4-GFP/UAS-mcherry-RNAi -VALIUM20 | 8 larvae/ 21 NMJs | Syt4-GFP intensity levels<br>A, B | Pre: Unpaired t-test, p<0.05<br>Post: Mann-Whitney, p<0.0001 |
|  | Gal4<sup>C380</sup>/Y;; Syt4-GFP/UAS-Tsg101-RNAi | 8 larvae/ 17 NMJs |  |  |
| **1B, F**<br>**SDCM** | Gal4<sup>Vglutx</sup>/Y; UAS-Evi-GFP/+; UAS-mcherry-RNAi-VALIUM20/+ | 7 larvae/ 21 NMJs | Evi-GFP intensity levels<br>A, B | Pre: Mann-Whitney, p<0.001<br>Post: Mann-Whitney, p<0.001 |
|  | Gal4<sup>Vglutx</sup>/Y; UAS-Evi-GFP/+; UAS-Tsg101-RNAi/+ | 8 larvae/ 25 NMJs |  |  |
| **1C, D, G, H**<br>**SDCM** | Gal4<sup>Vglutx</sup>/Y; UAS-APP-GFP/+; UAS-mcherry-RNAi-VALIUM20/+ | 6 larvae/ 15 NMJs | APP-GFP intensity levels<br>A, B | Pre: Mann-Whitney, p=0.56 n.s.<br>Post: Unpaired t-test, p<0.05 |
|  | Gal4<sup>Vglutx</sup>/Y; UAS-APP-GFP/+; UAS-Tsg101-RNAi/+ | 6 larvae/ 19 NMJs | Nrg intensity levels<br>A, B | Pre: Mann-Whitney, p=0.31 n.s.<br>Post: Mann-Whitney, p<0.0001 |
| **S1A**<br>**Airyscan** | Gal4<sup>Vglutx</sup>/w; UAS- Tsp42Ej-3xHA/+; UAS-mcherry-RNAi-VALIUM20/+ | 6 larvae/ 17 NMJs | No measurements |  |
|  | Gal4<sup>Vglutx</sup>/w; UAS- Tsp42Ej-3xHA/+; UAS-Tsg101-RNAi/+ | 6 larvae/ 19 NMJs<br>(Female) |  |  |
| **S1B-E**<br>**STED** | Gal4<sup>Vglutx</sup>/Y; UAS-APP-GFP/+; UAS-mcherry-RNAi-VALIUM20/+ | 6 larvae/ 19 NMJs | Quantification of density of APP-GFP and Nrg puncta | Nrg puncta density: 2-way ANOVA with Sidak's multiple comparison's test, p<0.001<br>APP puncta density: 2-way ANOVA with Sidak's multiple comparison's test, p<0.001 |
|  | Gal4<sup>Vglutx</sup>/Y; UAS-APP-GFP/+; UAS-Tsg101-RNAi/+ | 6 larvae/ 17 NMJs | APP-GFP puncta diameter | No statistical comparison made |
| **2A,D**<br>**SDCM** | Syt4-GFP<br>*Hrs<sup>D28</sup>/Df(2L)Exel6277*; Syt4-GFP<br>*nwk<sup>2</sup>*; Syt4GFP | 6 larvae /22 NMJs<br>7 larvae/ 24 NMJs<br>6 larvae/ 20 NMJs | Syt4-GFP intensity levels<br>A, B | Pre: One-way ANOVA with Tukey's multiple comparisons test<br>p<0.001<br>Post: Kruskal-Wallis with Dunn's multiple comparisons test<br>p<0.001 |
| **2B,E**<br>**SDCM** | Gal4<sup>Vglutx</sup>/Y; UAS-Evi-GFP/+<br>Gal4<sup>Vglutx</sup>/Y; UAS-Evi-GFP, *Hrs<sup>D28</sup>/Df(2L)Exel6277* | 7 larvae /24 NMJs<br>7 larvae/ 26 NMJs | Evi-GFP intensity levels<br>A, B | Pre: Mann-Whitney<br>p<0.001<br>Post: Mann-Whitney<br>p<0.001 |
| **2C,F**<br>**SDCM** | w<sup>1118</sup><br>*Hrs<sup>D28</sup>/Df(2L)Exel6277* | 8 larvae/ 24 NMJs<br>8 larvae/ 26 NMJs | Nrg intensity levels<br>A, B | Pre: Mann-Whitney, p<0.001<br>Post: Mann-Whitney, p<0.0001 |
| **2G-I**<br>**SDCM** | Gal4<sup>C380</sup>/Y; Syt4-GFP/UAS-mcherry-RNAi-VALIUM20<br>Gal4<sup>C380</sup>/Y; UAS-Shrub-RNAi/+ ; Syt4-GFP/+ | 6 larvae/ 22 NMJs<br>6 larvae/ 18 NMJs | Syt4-GFP intensity levels<br>A, B | Pre: Mann-Whitney, p=0.68 n.s.<br>Post: Mann-Whitney, p<0.001 |
|  |  |  | Nrg intensity levels<br>A, B | Pre: Mann-Whitney, p<0.01<br>Post: Mann-Whitney, p<0.001 |
| **2J-L**<br>**SDCM** | Gal4<sup>C380</sup>/Y;; Syt4-GFP/UAS-mcherry-RNAi- VALIUM20<br>Gal4<sup>C380</sup>/Y;; UAS-Vps4<sup>DN</sup>/+ ; Syt4-GFP/+ | 8 larvae/ 25 NMJs<br>8 larvae/ 27 NMJs<br>(Female) | Syt4-GFP intensity levels<br>A, B | Pre: Unpaired t-test, p<0.001<br>Post: Mann-Whitney, p<0.0001 |
|  |  |  | Nrg intensity levels<br>A, B | Pre: Mann-Whitney, p<0.0001<br>Post: Mann-Whitney, p<0.0001 |
| **3A, S2A**<br>**Airyscan** | Gal4<sup>Vglutx</sup>/Y; GFP-Rab5-KI/+<br>Gal4<sup>Vglutx</sup>/Y; GFP-Rab5-KI/+; UAS-Tsg101-RNAi/+ | 6 larvae/ 20 NMJs<br>6 larvae/ 17 NMJs | Mander's coefficients (fraction of Nrg in GFP-Rab5 positive puncta and fraction of GFP-Rab5 in Nrg positive puncta) | M1: Mann-Whitney, p<0.0001<br>M2: Unpaired t-test, p<0.0001 |
| **3A, S2B**<br>**Airyscan** | Gal4<sup>Vglutx</sup>/Y;; YFP-myc-Rab7KI/+<br>Gal4<sup>Vglutx</sup>/Y;; YFP-myc-Rab7KI, UAS-Tsg101-RNAi/+ | 7 larvae/ 20 NMJs<br>7 larvae/ 17 NMJs | Mander's coefficients (fraction of Nrg in YFP-Rab7 positive puncta and fraction of YFP-Rab7 in Nrg positive puncta) | M1: Mann-Whitney, p<0.01<br>M2: Unpaired t-test, p<0.01 |
| **3A, S2C**<br>**Airyscan** | Gal4<sup>C380</sup>/Y;; Syt4-GFP/UAS-mcherry-RNAi- VALIUM20<br>Gal4<sup>C380</sup>/Y;; Syt4GFP/UAS-Tsg101-RNAi | 7 larvae/ 25 NMJs<br>9 larvae/ 25 NMJs | Mander's coefficients (fraction of Syt4-GFP in anti-Rab11 positive puncta and fraction of anti-Rab11 in Syt4-GFP positive puncta) | M1: Unpaired t-test, p<0.0001<br>M2: Mann-Whitney: p<0.0001 |
| **3A, S2D-F**<br>**Rab11**<br>**Airyscan** | Gal4<sup>C380</sup>/Y;; Syt4-GFP/UAS-mcherry-RNAi- VALIUM20<br>Gal4<sup>C380</sup>/Y;; Syt4GFP/UAS-Tsg101-RNAi | 7 larvae/ 23 NMJs<br>9 larvae/ 25 NMJs | Rab11 intensity levels A | Unpaired t-test, p<0.001 |
|  |  |  | Number of anti-Rab11 positive puncta in HRP volume (puncta count/um<sup>3</sup>) | Mann-Whitney, P>0.05 ns |
|  |  |  | Average anti-Rab11 puncta size (total Rab11 volume um<sup>3</sup>/puncta number) | Mann-Whitney, p<0.05 |
| **3A, S2D-F Rab5**<br>**Airyscan** | Gal4<sup>Vglutx</sup>/Y; GFP-Rab5-KI/+<br>Gal4<sup>Vglutx</sup>/Y; GFP-Rab5-KI/+; UAS-Tsg101-RNAi/+ | 6 larvae/ 20 NMJs<br>6 larvae/ 17 NMJs | GFP-Rab5 intensity levels A | Mann-Whitney, p<0.05 |
|  |  |  | Number of GFP-Rab5 positive puncta in HRP volume (puncta count/um<sup>3</sup>) | Unpaired t-test, p<0.05 |
|  |  |  | Average GFP-Rab5 puncta size (total Rab5 volume um<sup>3</sup>/puncta number) | Unpaired t-test, p<0.001 |
| **3A, S2D-F Rab7**<br>**Airyscan** | Gal4<sup>Vglutx</sup>/Y;; YFP-myc-Rab7KI/+<br>Gal4<sup>Vglutx</sup>/Y;; YFP-myc-Rab7KI, UAS-Tsg101-RNAi/+ | 7 larva/ 20 NMJs<br>7 larva/ 17 NMJs | YFP-Rab7 intensity levels A | Mann-Whitney, p=0.14 n.s. |
|  |  |  | Number of YFP-Rab7 positive puncta in HRP volume (puncta count/um<sup>3</sup>) | Mann-Whitney, p=0.73 n.s. |
|  |  |  | Average YFP-Rab7 puncta size (total Rab7 volume um<sup>3</sup>/puncta number) | Mann-Whitney, p=0.25 n.s. |
| **3B**<br>**TEM** | Gal4<sup>Vglutx</sup>/Y;; UAS-mcherry-RNAi- VALIUM20/+<br>Gal4<sup>Vglutx</sup>/Y;; UAS-Tsg101RNAi/+ | 40 boutons from 3 larvae<br>56 boutons from 3 larvae | Fraction of images with three or more autophagic vacuoles | Fisher's exact test, p<0.0001 |
| **3C-D**<br>**SDCM** | Gal4<sup>Vglutx</sup>/Y;; UAS-mcherry-RNAi- VALIUM20/+ | 8 larvae/ 30 NMJs | Nrg intensity levels A, B | Pre: Unpaired t-test, p<0.01<br>Post: Mann-Whitney, p<0.05 |
|  | Gal4<sup>Vglutx</sup>/Y;; UAS-Atg1-RNAi/+ | 8 larvae/ 22 NMJs |  |  |
| **3E-G, S2J**<br>**Airyscan** | Gal4<sup>Vglutx</sup>/Y; UAS-GFP-mCherry-Atg8/+; UAS-Luciferase-RNAi-VALIUM10/+<br>Gal4<sup>Vglutx</sup>/Y; UAS-GFP-mCherry-Atg8/+; UAS-Tsg101-RNAi/+<br>Gal4<sup>Vglutx</sup>/Y; UAS-GFP-mCherry-Atg8/+<br>Gal4<sup>Vglutx</sup>/Y; UAS-GFP-mCherry-Atg8,<br>*Hrs*<sup>D28</sup>/Df(2L)Exel6277 | 8 larvae each<br>4 male, 4 female<br>per genotype | (2E-G) Fraction of grouped cell bodies volume occupied by GFP signal or mCherry signal. (S2D) Mander's coefficient for fraction total mCherry signal in GFP-positive volume | GFP in Tsg101: Mann-Whitney, p<0.001<br>mCherry in Tsg101: Mann-Whitney, p<0.001<br>GFP in Hrs: Mann-Whitney, p=0.35 n.s.<br>mCherry in Hrs: Unpaired t-test, p <0.001<br>Mander's Hrs: Mann-Whitney, p=0.3 n.s.<br>Mander's Tsg101: Mann-Whitney, p<0.001 |
| **S2G**<br>**SDCM** | Gal4<sup>Vglutx</sup>/Y; UAS-GFP-mCherry-Atg8/+; UAS-Luciferase-RNAi-VALIUM10/+<br>Gal4<sup>Vglutx</sup>/Y; UAS-GFP-mCherry-Atg8/+; UAS-Atg1-RNAi/+ | 2 larvae<br>3 larvae | No measurements | NA |
| **S2H-I**<br>**SDCM** | w<sup>1118</sup><br>*Atg2*<sup>EP3697</sup>/Df(3L)Exel6091 | 8 larvae/ 18 NMJs<br>7 larvae/ 17 NMJs<br>Mixed males and females | Nrg intensity levels<br>A, B | Pre: Mann-Whitney, p<0.01<br>Post: Unpaired t-test, p<0.001 |
| **4A**<br>**SDCM** | Gal4<sup>C380</sup>/Y;; Syt4-GFP/UAS-mcherryRNAi-VALIUM20<br>Gal4<sup>C380</sup>/Y;; Syt4-GFP/UAS-Tsg101RNAi | 10 larvae<br>8 larvae | Intensity measurement from an ROI containing 6-8 cell bodies in ImageJ | Unpaired t-test, p<0.0001 |
| **4B**<br>**SDCM** | Gal4<sup>C380</sup>/Y;; Syt4-GFP/UAS-mcherryRNAi- VALIUM20<br>Gal4<sup>C380</sup>/Y;; Syt4-GFP/UAS-Tsg101RNAi | 10 larvae<br>9 larvae | Intensity measurement from HRP thresholded axon bundles proximal (within ~100-300µm) to the ventral ganglion in ImageJ | Unpaired t-test, p<0.05 |
| **4C-E**<br>**SDCM** | Gal4<sup>Vglutx</sup>/Y; UAS-APPGFP/+; UAS-mcherryRNAi-VALIUM20/+<br>Gal4<sup>Vglutx</sup>/Y; UAS-APPGFP/+; UAS-Tsg101RNAi/+ | 12 larvae<br>12 larvae | Kymograph analysis, manual count of anterograde, retrograde, stationary, and complex moving APP-GFP particles in ImageJ | Unpaired t-test, Stationary: p<0.001<br>Retrograde: p0.48 n.s.<br>Anterograde: p=0.50<br>Complex: p=0.12 |
|  |  |  | Kymograph analysis, calculation of the slope of particle tracks, including segments from complex tracks in ImageJ | Retrograde: Mann-Whitney, p<0.05<br>Anterograde: Unpaired t-test p=0.68 n.s. |
| **S3A-D**<br>**SDCM** | Gal4<sup>Vglutx</sup>/Y; UAS-Mito-GFP/+; UAS-mcherry-RNAi-VALIUM20/+<br>Gal4<sup>Vglutx</sup>/Y; UAS-Mito-GFP/+; UAS-Tsg101-RNAi/+ | 11 larvae | Kymograph analysis, manual count of percentage of anterograde and retrograde mito-GFP particles in ImageJ | Retrograde: Unpaired t-test, p=0.70 n.s.<br>Anterograde: Unpaired t-test, p=0.70 n.s. |
|  |  | 11 larvae |  |  |
|  |  |  | Kymograph analysis, calculation of the slope of particle tracks in ImageJ | Retrograde: Unpaired t-test, p=0.65 n.s.<br>Anterograde: Unpaired t-test, p=0.36 n.s. |
|  |  |  | Intensity measurement from first frame of videos of axon bundles proximal (within ~100-300µm) to the ventral ganglion in ImageJ | Unpaired t-test, p=0.79 n.s. |
| **5A-B**<br>**SDCM** | Gal4<sup>Vglutx/Y;;</sup> UAS-mcherryRNAi- VALIUM20/+<br>Gal4<sup>Vglutx/Y;;</sup> UAS-Tsg101RNAi/+<br>*evi<sup>2</sup>/evi<sup>2</sup>* | 7 larvae/14 A2 NMJs, 12 A3 NMJs | Active zone number on muscle 6/7 | A2: One-way ANOVA with Tukey's multiple comparison's test, p<0.001<br>A3: One-way ANOVA with Tukey's multiple comparison's test, p<0.001 |
|  |  | 7 larvae/11 A2 NMJs, 12 A3 NMJs |  |  |
|  |  | 7 larvae/8 A2 NMJs, 12 A3 NMJs | Number of active zones per NMJ area on muscle 6/7 | A2: Kruskal-Wallis with Dunn's multiple comparison's test, p<0.001<br>A3: One-way ANOVA with Tukey's multiple comparison's test, p<0.001 |
|  |  |  | Number of boutons on muscle 6/7 | A2: One-way ANOVA with Tukey's multiple comparisons test, p<0.001<br>A3: One-way ANOVA with Tukey's multiple comparisons test, p<0.001 |
| **5C-E**<br>**SDCM** | Gal4<sup>Vglutx/Y;;</sup> UAS-mcherryRNAi- VALIUM20/+<br>Gal4<sup>Vglutx/Y;;</sup> UAS-Tsg101RNAi/+<br>*evi<sup>2</sup>/evi<sup>2</sup>* | 26 larvae/ 87 NMJs | Fraction of images displaying expanded "feathery" DLG pattern | Fishers exact test<br>WT vs *evi<sup>2</sup>*<br>p<0.0001<br>WT vs Tsg101<sup>KD</sup><br>p=0.27 n.s. |
|  |  | 26 larvae/ 87 NMJs |  |  |
|  |  | 29 larvae/ 91 NMJs |  |  |
|  |  | (Pooled data from 3 independent experiments, mix of males and females) | Ghost bouton number on muscle 6/7 (HRP-positive bouton with no postsynaptic Dlg) | A2: Kruskal-Wallis test with Dunn's multiple comparison's test, p<0.001<br>A3: Kruskal-Wallis test with Dunn's multiple comparison's test, p<0.05 |
|  | *Hrs<sup>D28</sup>/Df(2L)Exel6277*<br>Gal4<sup>Vglutx/Y;;</sup> UAS-Tsg101RNAi/+ | 12 larvae/40 NMJs<br>10 larvae/32 NMJs | Ghost bouton number on muscle 6/7 (HRP-positive bouton with no postsynaptic Dlg) | A2: Mann-Whitney test, p=0.005<br>A3: Mann-Whitney test, p=0.68 n.s. |
| **5F-G LSCM** | *y<sup>1</sup>, w<sup>1118</sup>*<br>Gal4<sup>Vglutx/Y;;</sup> UAS-Tsg101RNAi/+<br>*Hrs<sup>D28</sup>/Df(2L)Exel6277* | 4 larvae, 16 muscles, 264 nuclei<br>4 larvae, 16 muscles, 303 nuclei<br>4 larvae, 16 muscles, 274 nuclei | Fz2-C puncta number per nucleus (muscles 6 and 7) | One-way ANOVA with Tukey's multiple comparisons test, p<0.001 |
| **6A, B SDCM** | Gal4<sup>C380/Y;;</sup> Syt4GFP/UAS-mcherry-RNAi-VALIUM20<br>Gal4<sup>C380/Y;;</sup> Syt4-GFP/UAS-Tsg101RNAi | WT Mock: 6 larvae/ 22 NMJs<br>WT Stim: 5 larvae/ 23 NMJs<br>Tsg101 Mock: 6 larvae/ 25 NMJs<br>Tsg101 Stim: 5 larvae/ 23 NMJs | Ghost bouton number on muscle 4 (HRP-positive bouton with no postsynaptic Dlg) | WT stim vs mock: Mann-Whitney, p<0.001<br>Tsg101 stim vs mock: Mann-Whitney, p<0.001<br>WT stim vs Tsg101 stim: Mann-Whitney, p=0.81 n.s. |
| **6C, D electro-physiology** | Gal4<sup>Vglutx/+;;</sup> UAS-mcherry-RNAi-VALIUM20/+<br>Gal4<sup>Vglutx/+;;</sup> UAS-Tsg101RNAi/+ | 9 larvae / 12 NMJs<br>8 larvae / 12 NMJs | Fold change in mEJP frequency | No statistical measurement |
| **6E, F electro-physiology** | *w<sup>1118</sup>*<br>*Hrs<sup>D28</sup>/Df(2L)Exel6277*<br>*syt4<sup>BA1</sup>* | 9 larvae / 11 NMJs<br>12 larvae/12 NMJs<br>10 larvae/10 NMJs | Fold change in mEJP frequency | No statistical measurement |
| **S4B SDCM** | Gal4<sup>C380/Y or +;;</sup> Syt4<sup>TSTEP</sup><br>Gal4<sup>C380/Y or +;;</sup> Syt4<sup>TSTEP</sup>/UAS-Rippase, Syt4<sup>TSTEP</sup><br>*w;;* Gal4<sup>C57</sup>, Syt4<sup>TSTEP</sup>/UAS-Rippase, Syt4<sup>TSTEP</sup> | No quantification | No measurements | NA |
| **S4C, D LSCM** | Gal4<sup>Vglutx/+;;</sup> UAS-mcherry-RNAi-VALIUM20/+<br>Gal4<sup>Vglutx/+;;</sup> UAS-Tsg101-RNAi/+ | 7 larvae / 13 NMJs<br>7 larvae / 13 NMJs | Ghost bouton number on muscle 6/7 (HRP-positive bouton with no postsynaptic Dlg) | Unpaired t-test, p=0.65 n.s. |
| **7B, D SDCM** | Gal4<sup>C380/Y;</sup> Syt4-GFP<br>Gal4<sup>C380/Y;</sup> UAS-DegradGFP, Syt4-GFP | 29 NMJ branches<br>25 NMJ branches | Syt4-GFP intensity levels<br>A, B | Pre: Unpaired t-test, p<0.001<br>Post: Unpaired t-test, p<0.001 |
| **7C, E SDCM** | Syt4-GFP<br>UAS-DegradGFP; Gal4<sup>C57</sup>, Syt4-GFP | 8 larvae/ 33 NMJ branches<br>8 larvae/ 30 NMJ branches | Syt4-GFP intensity levels<br>A, B | Pre: Mann-Whitney, p=0.59 n.s.<br>Post: Mann-Whitney, p=0.56 n.s. |
| **7F**<br>**SDCM** | Genotypes from 7B and 7C | N from 4F and 4G | Number of Syt4-GFP positive puncta in HRP volume (puncta count/$\mu\text{m}^3$) | C380: Unpaired t-test, $p < 0.001$<br>C57: Mann-Whitney, $p = 0.84$ n.s. |
| **7G-H**<br>**SDCM** | UAS-DraperRNAi/+; Syt4-GFP/+<br>GAL4<sup>C155</sup>; UAS-DraperRNAi/+ Syt4-GFP<br>UAS-DraperRNAi/+ Syt4-GFP/GAL4<sup>C57</sup><br>UAS-DraperRNAi/+ Syt4-GFP/Gal4<sup>repo</sup><br>UAS-DraperRNAi/+ Syt4-GFP/GAL4<sup>tub</sup> | 6 larvae/ 26 NMJs<br>6 larvae/ 18 NMJs<br>6 larvae/ 26 NMJs<br>6 larvae/ 25 NMJs<br>6 larvae/ 28 NMJs | Syt4-GFP intensity levels<br>A, B | Pre: Kruskal-Wallis $P < 0.001$<br>Post: Kruskal-Wallis $P < 0.001$ |
| **S5A**<br>**SDCM** | Dlg1-MiMIC/+; CyOGFP/+;<br>Gal4<sup>C57</sup>, Syt4-GFP/TM6<br>Dlg1-MiMIC/+; UAS-DegradGFP/+; Gal4<sup>C57</sup>, Syt4-GFP/ Gal4<sup>C57</sup> | 4 larvae/ 14 NMJs<br>4 larvae/ 13 NMJs | No measurements | NA |
| **S5B-C**<br>**SDCM** | w<sup>1118</sup><br>GAL4<sup>C155</sup>; UAS-DraperRNAi/+<br>UAS-DraperRNAi/+ GAL4<sup>C57</sup>/+<br>UAS-DraperRNAi/+ GAL4<sup>repo</sup>/+ | 6 larvae/ 27 NMJs<br>6 larvae/ 25 NMJs<br>6 larvae/ 29 NMJs<br>6 larvae/ 31 NMJs | Draper mean intensity | NMJ: One-way ANOVA $p < 0.001$<br>Axon: Kruskal-Wallis $p < 0.001$ |

